# PepFlow: direct conformational sampling from peptide energy landscapes through hypernetwork-conditioned diffusion

**DOI:** 10.1101/2023.06.25.546443

**Authors:** Osama Abdin, Philip M. Kim

**Affiliations:** Department of Molecular Genetics, University of Toronto, Toronto, ON M5S 3E1, Canada; Department of Computer Science, University of Toronto, Toronto, ON M5S 3E1, Canada; Donnelly Centre for Cellular and Biomolecular Research, University of Toronto, Toronto, ON M5S 3E1, Canada

## Abstract

Deep learning approaches have spurred substantial advances in the single-state prediction of biomolecular structures. The function of biomolecules is, however, dependent on the range of conformations they can assume. This is especially true for peptides, a highly flexible class of molecules that are involved in numerous biological processes and are of high interest as therapeutics. Here, we introduce PepFlow, a generalized Boltzmann generator that enables direct all-atom sampling from the allowable conformational space of input peptides. We train the model in a diffusion framework and subsequently use an equivalent flow to perform conformational sampling. To overcome the prohibitive cost of generalized all-atom modelling, we modularize the generation process and integrate a hyper-network to predict sequence-specific network parameters. PepFlow accurately predicts peptide structures and effectively recapitulates experimental peptide ensembles at a fraction of the running time of traditional approaches. PepFlow can additionally be used to sample conformations that satisfy constraints such as macrocyclization.

## 1 Introduction

Protein-peptide interactions are ubiquitous across molecular pathways and are integral to many cellular functions. In total, it’s estimated that up to 40% of protein-protein interactions are mediated by peptide binding [1]. These interactions involve the binding of globular proteins to short segments that are typically localized to disordered regions [2]. Outside of their endogenous function, short peptides also have several properties that make them amenable to therapeutic development. Compared to small molecules, peptides tend to be more specific, and present a lower risk for toxicity [3]. Compared to larger biologics, peptides are cheaper to produce and tend to be less immunogenic [3]. Overall, peptide therapeutics contribute to a substantial share of the pharmaceutical market. Glucagon-like peptide 1 (GLP1) analogue peptides, for example, are consistently top selling drugs [3]. Given their prevalence in biological processes and potential for therapeutic application, there is a need for computational tools to expedite the modelling and engineering of peptides.

In general, deep learning has driven substantial improvements in biomolecular modelling, with the clearest example being the unprecedented success of AlphaFold2 (AF2) for the single-state modelling of protein structures [4]. Despite being trained on proteins, AF2 has improved on the state-of-the-art for modelling the structures of linear peptides [5], cyclic peptides [6], and peptide-protein complexes [7]. Nevertheless, AF2 exhibits a number of failure cases and does not capture the range of conformations a peptide can assume. Modelling the flexibility of peptides is crucial to attaining a complete understanding of their function and binding.

There have been attempts to extend AF2 for conformational sampling by exploiting inherent stochasticity in the structure prediction pipeline. One approach involves sub-sampling the multiple sequence alignments (MSAs) used in structure prediction to generate different structural conformations [8]. Another approach makes use of the dropout layers in the network to induce variability in the output [9]. While these alterations to AF2 do succeed in generating different structural conformations, they do not predict the propensity of the proteins to fold into the different structures, and do not necessarily model the underlying energy landscape.

In addition to methods that make use of the pre-trained AF2 network, there has also been work on generative models that can be used to directly sample protein conformations. Recently, a deep learning method was developed that equips the embeddings from a single-structure prediction approach [10] with a diffusion model to directly generate different structures [11]. While this model has some capacity for predicting alternate conformations, it was shown to perform better as a single-structure predictor. Another method was developed that uses a generative adversarial network (GAN) trained on molecular dynamics (MD) simulations to generate ensembles of intrinsically disordered proteins, and was shown to be able to do so with considerable accuracy [12].

Boltzmann generators represent an alternative paradigm in using deep learning for conformational sampling [13]. Boltzmann generators overcome the limitations of trajectory-based statistical mechanics approaches by using deep learning models for direct conformational sampling. More specifically, they make use of normalizing flows to learn the distribution of conformations for a particular molecular system. Compared to other generative models, normalizing flows have the advantage of enabling exact likelihood computation [14]. This in turn allows for training by energy, a process by which the model is optimized to match the distribution of states given by a particular molecular force field. Given a function, *u*(**x**), that defines the energy of a particular configuration of the molecular system, the equilibrium distribution of states would equal the Boltzmann distribution, which is proportional to *e^−u^*^(^*^x^*^)^.

Typically, Boltzmann generators have been restricted to modelling single molecular systems. A recent study extended this work, however, to develop a generalized Boltzmann generator that can be used to model the distribution of torionsal angles for different small molecules [15]. The authors leveraged score-based generative models (SGMs) to develop a method that can be used to effectively sample molecular conformers. Score-based generative models are a powerful class of diffusion models that use stochastic differential equations (SDEs) to gradually noise data to a simple prior distribution [16]. Although SGMs do not directly permit exact likelihood computation, it has been shown that for each process specified by an SDE, there exists an equivalent ordinary differential equation (ODE) that specifies the same marginal probability distribution at each time step [16]. This probability flow ODE can be used for exact likelihood computation [16], and this approach was successfully used to perform energy-based training with the aforementioned generalized Boltzmann generator [15].

In this work, we sought to develop an approach that could be used for the direct, all-atom sampling of peptide conformations. Even for short peptides, performing all-atom sampling that is both accurate and efficient poses a substantial challenge. A peptide of length 15 contains up to 377 atoms and 1131 degrees of freedom within its structure. Without *a priori* knowledge of the molecule topology, modelling the inter-dependencies between degrees of freedom for an arbitrary peptide sequence is costly. To address this, we developed PepFlow, a modular, hypernetwork-conditioned Boltzmann generator that generates all-atom conformations for any input peptide sequence.

To train this model, we exploit the diversity of fragment conformations found in the Protein Data Bank (PDB) as an initial training source, before fine-tuning on peptide conformations and training by energy. PepFlow is trained on known molecular conformations as an SGM and is subsequently used as a probability flow ODE for sampling and training by energy. We show that PepFlow has a large capacity to predict both single-state structures and conformational ensembles. In particular, PepFlow can recapitulate structures found in experimentally generated ensembles of short linear motifs (SLiMs). Learning the conformational landscape of peptides additionally allows the modelling of therapeutically relevant peptide modifications, such as macrocyclization, through latent space conformational searches.

## 2 Results

### 2.1 PepFlow recapitulates molecular features of peptides

The PepFlow architecture has two defining features; PepFlow is separated into three networks that progressively model the peptide conformation, and PepFlow makes use of a hypernetwork to generate a subset of sequence-specific parameters (Figure 1a). The first of the three networks is used to model the backbone atoms and the centroid of the side chain atoms, implicitly capturing most of the flexibility of the input peptide. Input peptide coordinates are represented as a fully connected graph and passed through a set of E(3) Equivariant Graph Neural Network (EGNN) layers [17] that have sequence-specific parameters predicted by an attention-based hypernetwork, and a set of general EGNN layers that are directly optimized (Figure 1a). Use of a hypernetwork allows for increased model expressitivity without a commensurate increase in the size of the dynamics network.

**Figure 1:**
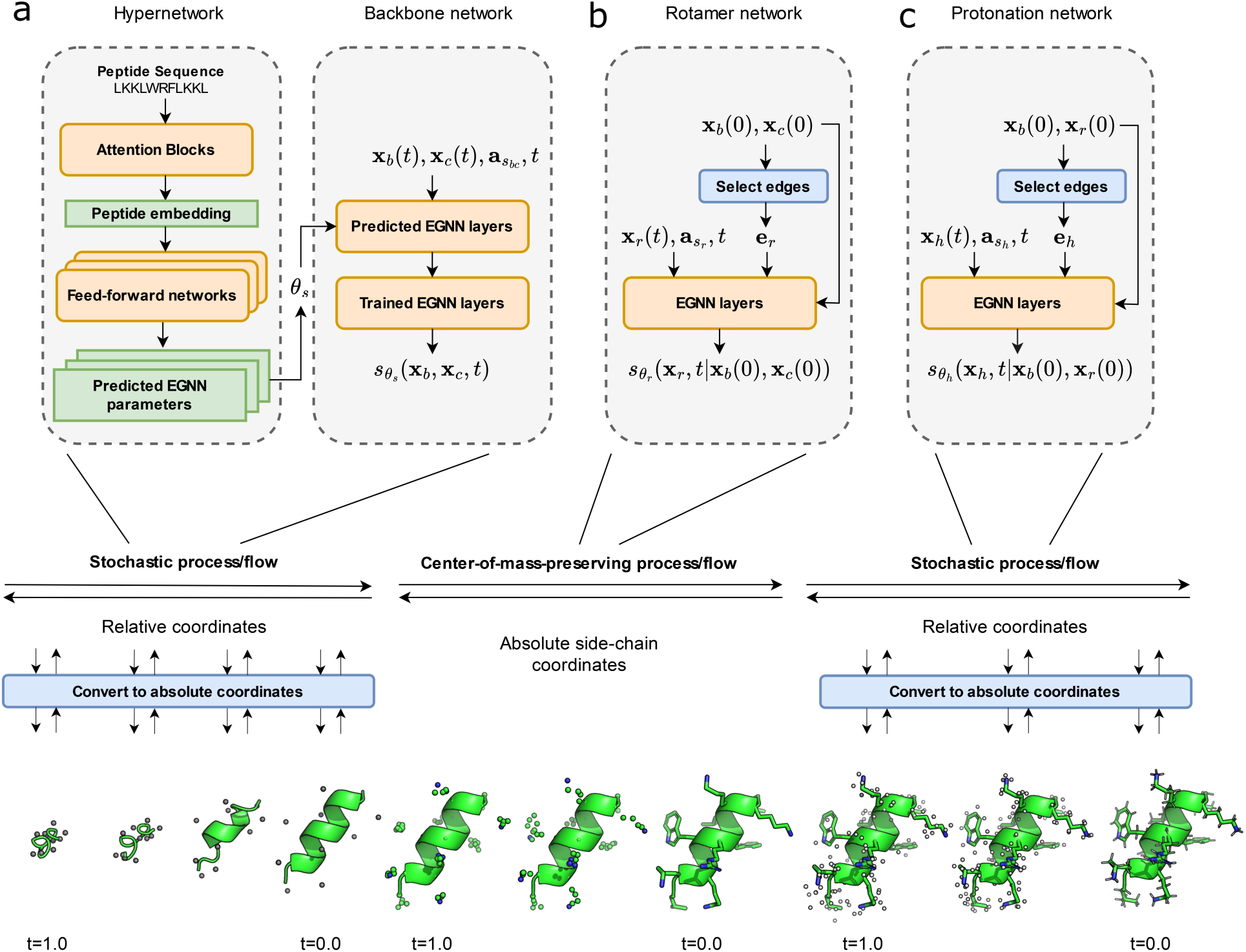
Schematic of the PepFlow architecture a. A hypernetwork generates a subset of sequence-specific network parameters that are used by an EGNN-based score network to facilitate backbone generation. b. An EGNN-based score network is used with a CoM preserving process to generate side chain heavy atom coordinates while enforcing side chain centroids. c. An EGNN-based score network is used to protonate peptides based on input heavy atom coordinates with edges determined by heavy atom pairwise distances.

Notably, rather than denoising the absolute coordinates of the atoms, the model learns to denoise relative coordinates between each atom and its N-terminal neighbor, with side chain centroid positions encoded relative to the C*α* atom. While the network layers still operate on absolute coordinates, this changes the noising process so that it gradually moves the peptide backbone to an ideal chain (Supplementary Information). Similar to changes applied to previous generative protein models, this modification integrates an inductive bias into the model that reflects the peptide’s properties as a chain [11, 18].

After training this model on protein fragments from the PDB, we assessed its ability to generate reasonable molecules given input protein fragment sequences. Sampling was done using the ODE flow (see Methods) and despite the fact that samples generated this way tend to be lower quality than those generated using predictor-corrector samplers [16], we found that they accurately capture peptide molecular characteristics. More specifically, the bond lengths, bond angles, and torsional angles were all in line with experimentally-solved structures (Figure S1a-c). The distributions of side chain centroid positions were also found to match the ground truth data (Figure S1d).

The second network is used to model the heavy atoms in the side chain based on the generated backbone and centroids (Figure 1b). Unlike the backbone model, which operates on fully connected graphs, edges were restricted to residues within 8 Å of each other’s side chain centroids. To allow tractable likelihood computation over the *real* peptide atomic coordinates, the model enforces the condition that *p*_0_(**x***_b_*(0), **x***_r_*(0)) = *p*_0_(**x***_b_*(0), **x***_r_*(0), **x***_c_*(0)), where **x***_b_*(0), **x***_r_*(0) and **x***_c_*(0) are the backbone coordinates, rotamer coordinates and side chain centroid coordinates at time 0, respectively. In practice, this is done by fixing the centroids of the generated side chains to the input centroid through a center-of-mass (CoM) preserving process (Supplementary Information).

We find that the resulting rotamer model effectively generates side chain conformations given backbone and centroid information, with consistently low root-mean square deviations (RMSDs) on the fragment validation set (Figure S2). Performing latent space temperature-scaling further lowers the RMSDs of generated side chains (Figure S2b). As expected, generated conformations for residues with more atoms and rotatable bonds were found to have higher RMSDs on average (Figure S2c). We additionally found that the distribution of *χ* angles for the different residues does closely match the empirical distribution, with the main discrepancies being due to ambiguous atom labelling at the end of a side chain (Figure S3). This suggests that the model generates reasonable conformations even when they deviate from the ground-truth positions.

The final network is used to model the positions of protons based on the coordinates of heavy-atoms in the peptide (Figure 1c). More specifically, the model denoises the relative positions of hydrogen atoms to their bound heavy atom. The processed graph contains two sets of edges. The first set contains edges between hydrogen atoms that are covalently linked to the same heavy atom. The second set contains edges between hydrogen atoms and heavy atoms within 4 Å of their bound atom. Unlike the backbone and rotamer models which were trained on fragments from crystal structures in the PDB, this model was trained directly on a subset of molecular dynamics (MD) simulations from the Database of Antimicrobial Activity and Structure of Peptides (DBAASP) [19] due to the absence of hydrogens from most experimentally solved structures. After training this model, we tested its ability to re-protonate peptide conformations derived from a set of MD simulations. We found that the generated conformations matched closely with the ground-truth MDs, with highly correlated energies and hydrogen bond counts (Figure S4).

### 2.2 PepFlow effectively generates sequence-specific peptide conformations

After initial training on known molecular conformations, PepFlow is able to generate a variety of all-atom conformations for peptides of different lengths (Figure S5). We next assessed the ability of PepFlow to generate sequence-specific conformations, and the extent to which this is mediated by the hypernetwork. Different models were trained after varying the number of layers with parameters predicted by the hypernetwork. Using these models, conformations were sampled for different fragment sequences in the validation set. We found that predictions for helical and coiled fragments showed minimal variation with the different models. Increasing the number of hypernetwork-predicted layers resulted in predictions that were, on average, slightly more structured (Figure S6a,b). A much more noticeable difference is seen when predicting conformations for strand-forming fragments. Increasing the number of parameters predicted by the hypernetwork drastically increases the propensity of PepFlow to generate peptides forming strands (Figure S6a,b), suggesting that that the hypernetwork plays an important role in predicting longer range interactions.

The model was further fine-tuned on solved peptide structures. The fine-tuned model shows a substantial increase in the number of predicted coiled conformations relative to the number of predicted structured conformations (Figure S7a), consistent with the flexible nature of peptides. The fine-tuned model additionally predicts solved peptide structural conformations with a much higher degree of accuracy than the model trained solely on fragment conformations (Figure S7b,c).

Using this finetuned model, we evaluated the ability of PepFlow to predict conformations of peptides from a test set of recent, non-redundant, structures from the PDB. To generate a single-structure prediction, 100 conformations were sampled with PepFlow, the conformations were clustered, and the centroid of the cluster with the most members was used as the prediction (see Methods). In most cases, the size of derived clusters was found to be negatively correlated with C*α* RMSD (Figure S8). We compared the performance of PepFlow to AF2, the current state-of-the-art in peptide structure prediction [5], and ESM-Fold [20], a single-sequence structure prediction method. PepFlow performs comparably to both these methods, and outperforms them in many cases (Figure 2a-c). This includes cases where PepFlow captures subtle features of solved structures as well as cases where PepFlow correctly predicts conformations that AF2 and ESMFold incorrectly model (Figure 2d). Interestingly, PepFlow and AF2 differ in terms of the cases on which they perform well (Figure 2b), indicating that the models are somewhat orthogonal.

**Figure 2:**
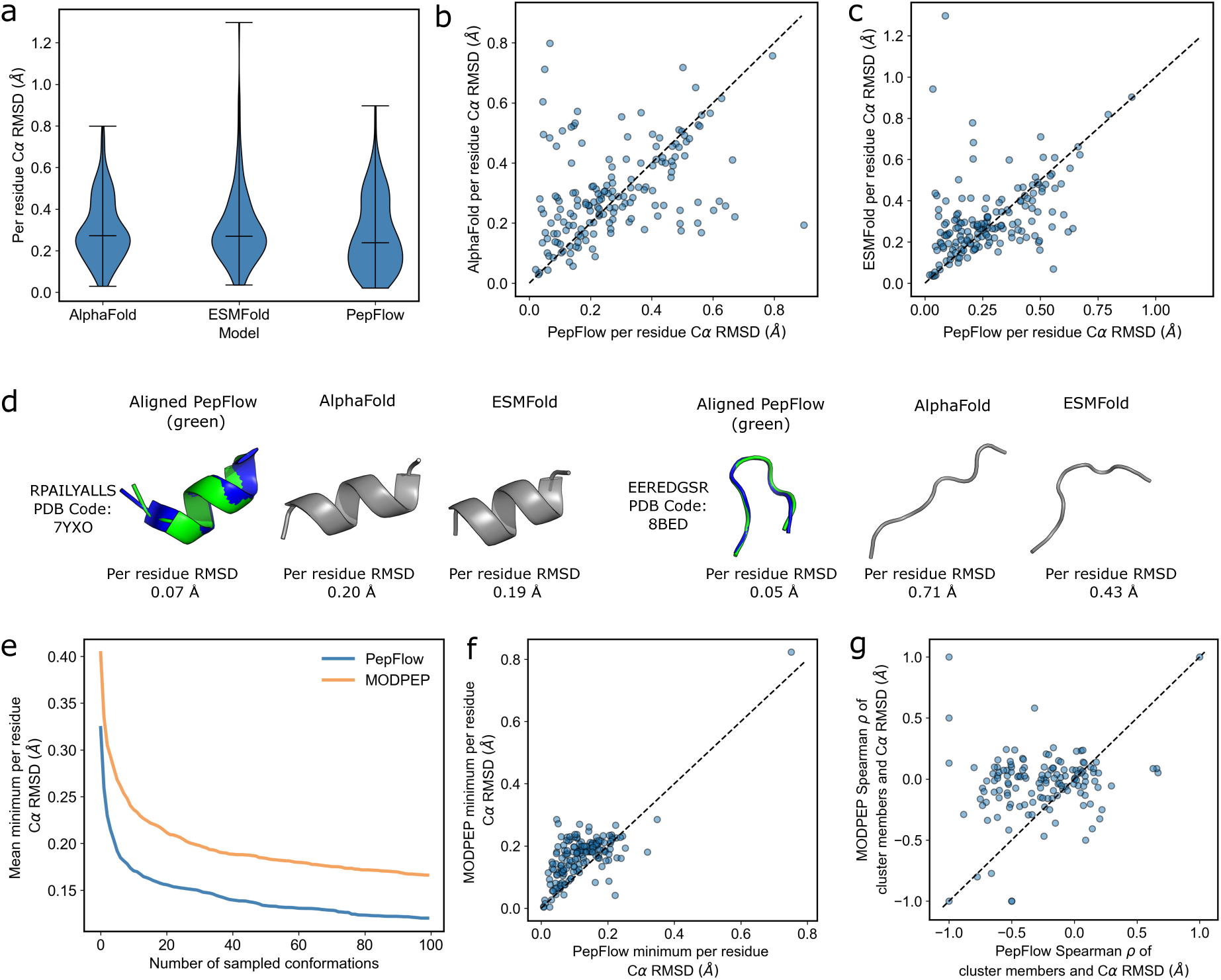
PepFlow prediction of experimentally solved structures. a. Distribution of C*α* RMSDs to peptide test set with PepFlow, AF2 and ESMFold. b. Comparison of PepFlow predictions on a test set of solved peptide structures with AF2. c. Comparison of PepFlow predictions on a test set of solved peptide structures with ESMFold. d. Example peptides where PepFlow outperforms AF2 and ESMFold. e. Minimum RMSD of generated samples from PepFlow and MODPEP as more conformations are generated. f. Comparison of minimum RMSDs to solved peptide structures with 100 conformations generated using PepFlow and MODPEP. g. Comparison of correlations of cluster size to C*α* RMSD using PepFlow and MODPEP.

It is important to note that a majority of peptide structures in the PDB are solved as a complex with a globular protein. Given the fact that peptides often undergo conformational changes upon binding [21], single-structure prediction in the absence of this context is necessarily going to be lacking. We therefore assessed whether PepFlow can capture bound peptide conformations as more samples are generated. As a baseline, PepFlow was compared to the MODPEP method [22], which generates conformations through secondary structure prediction and assembly of fragments from known bound peptide conformations. We find that PepFlow outperforms MODPEP in the vast majority of cases, and can accurately capture the conformations of solved peptide structures (Figure 2e,f). We additionally ranked the conformations generated by PepFlow and MODPEP using the aforementioned clustering approach, and found stronger correlations with PepFlow-generated conformations (Figure 2g), indicating that PepFlow more frequently generates accurate peptide structures.

### 2.3 Training by energy enables peptide conformational ensemble prediction

After finetuning PepFlow on peptide structures, training by energy was performed by sampling peptide conformations for different sequences, and minimizing the Kullback-Leibler (KL) divergence between the model proposal distribution and the unnormalized Boltzmann distribution parameterized by a molecular force field (see Methods). It has been previously shown that a Boltzmann generator trained only with this objective will learn to sample from a single metastable state [13]. We therefore added an auxiliary score-matching loss using MD-generated conformations from a subset of the DBAASP dataset.

To assess the effect of training by energy on ensemble generation, we sampled 100 conformations with PepFlow before and after training by energy for sequences in the MD validation set, with latent sampling scaled by different temperatures. These conformations were compared to the MD-derived ensemble, consisting of 200 snapshots taken every 2 ns in a 400 ns simulation. Overall, PepFlow-generated ensembles after training by energy were found to resemble the MD ensembles more closely, and performance was found to improve with latent temperature scaling. In particular, the distribution of radius of gyration (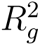) and the distribution of pairwise C*α* distances were found to match the MD-derived ensemble more closely after training by energy (Figure S9a, b). We also found that PepFlow conformations captured a much larger percentage of the conformations generated in the MD simulations, and that a much larger percentage of PepFlow conformations were represented in the MD ensemble after training by energy (Figure 3a,b, S9c,d). We additionally evaluated the impact of training by energy on the rotamer and protonation networks. Rotamer generation showed a similar degree of agreement with the MD ensembles after training by energy (Figure S10), but training by energy was found to drastically improve peptide protonation (Figure S11).

**Figure 3:**
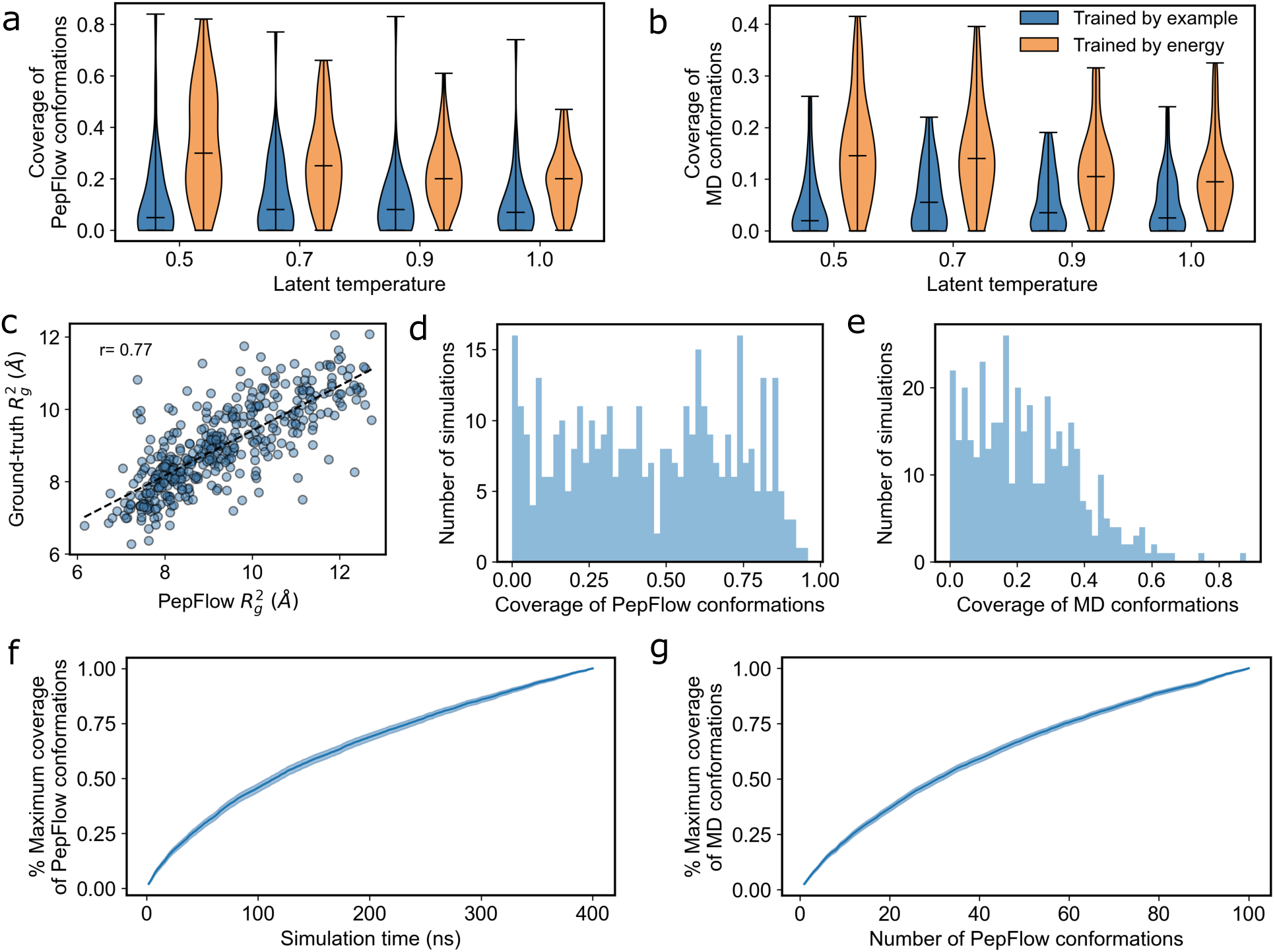
Comparison of PepFlow-generated ensembles to MD simulations a. Coverage of PepFlow-sampled conformations by validation MD conformations before and after training by energy at different latent temperatures. b. Coverage of validation MD conformations by PepFlow-sampled conformations before and after training by energy at different latent temperatures. c. Comparison of 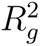 between PepFlow and test MD conformations. d. Distribution of coverage of PepFlow-sampled conformations by test MD conformations. e. Distribution of coverage of test MD conformations by PepFlow-sampled conformations. f. Coverage of PepFlow-generated conformations with simulation time, 90% confidence interval is shown. b. Coverage of MD conformations with increased PepFlow sampling, 90% confidence interval is shown.

To evaluate the generalizability of this performance, we similarly benchmarked PepFlow on a test set of 401 non-redundant MD simulations, and observed similar performance. In particular, we found that the mean 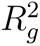 of predicted conformations is well correlated with the 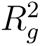 of MD conformations (Figure 3c). On average, at a threshold of 0.15 Å C*α* RMSD per residue, we found that 22.8% of MD conformations were covered by 100 PepFlow samples, and 44.6% of PepFlow conformations were covered by 400 ns of MD (Figure 3d,e). Notably, coverage of both sets of conformations is continually increasing with additional sampling, likely a product of the flexibility of the peptides being modelled (Figure 3f, g).

A more granular analysis of generated conformations shows that even at low coverage, the overall distribution of PepFlow-generated conformations can be highly concordant with the results of the MD. Analysis of peptides with varying secondary structure propensities shows that PepFlow sampling can reproduce a variety of different peptide structures (Figure S12-14). Importantly, PepFlow generates disparate modes that are represented by different regions of the MD ensemble (Figure S12-14). Projecting the generated PepFlow coordinates onto principal components fit to the MD ensemble shows that despite only sampling a small number of conformations, PepFlow captures a significant amount of the variation seen in the MD simulations (Figure S12-14).

### 2.4 PepFlow can efficiently recapitulate experimental ensembles

We next assessed the ability of PepFlow to predict experimental ensembles. To this end, we made use of the Protein Ensemble Database (PED), a curated database of structural ensembles generated using various experimental methods [23]. This database includes a number of intrinsically disordered proteins, which tend to be enriched in SLiMs. Using the eukaryotic linear motif (ELM) webserver [24], we annotated SLiMs of length 8-15 amino acids in the proteins composing the PED. The derived SLiMs show various degrees of disorder, with some occurring in highly structured regions and others occurring in regions of high disorder (Figure S15).

To determine the degree to which different annotated SLiMs display peptide-like characteristics, we determined the tertiary contacts found in each fragment based on Cartesian distances to amino acids that are distant in primary sequence (see Methods). For each of the SLiMs in the dataset, we sampled 100 PepFlow conformations and as expected, we found that the performance of PepFlow is negatively correlated with the number of tertiary contacts in the peptide (Figure S16). We more extensively validated PepFlow on peptide sequences with fewer than the lenient threshold of three tertiary contacts per residue. For each of these remaining sequences, 1000 conformations were sampled using PepFlow. Overall, the 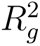 of generated samples is well-correlated with the experimental ensembles (Figure 4a). Additionally, we found that on average 46.3% of PepFlow conformations are covered by the experimental ensembles at a threshold of 0.15 Å RMSD per residue (Figure 4b). This coverage is highly correlated with the total number of experimental models of the SLiMs and begins to converge at around 1e3 conformations (Figure S17a,b). We furthermore found that an average of 46.5% of the SLiM conformations are covered by PepFlow samples, and we observe more than 30% coverage in 84% of the samples (Figure 4c). This value was also found to continually increase with additional PepFlow sampling (Figure S17c).

**Figure 4:**
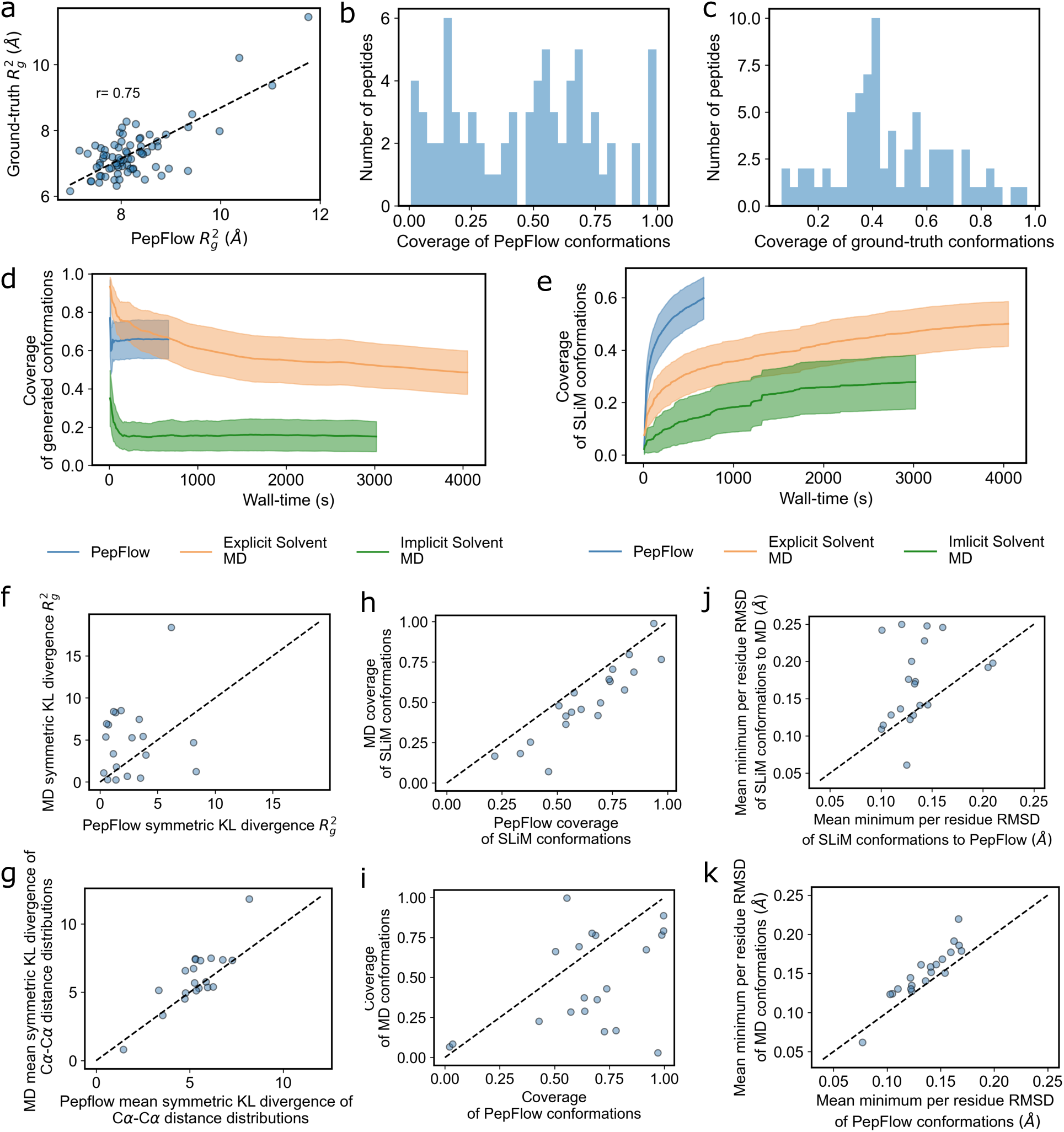
Performance of PepFlow and MD simulations on SLiM ensemble generation a. Comparison of 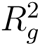 between PepFlow and ground-truth conformations. b. Distribution of coverage of PepFlow-sampled conformations by ground-truth conformations. c. Distribution of coverage of ground-truth conformations by PepFlow-sampled conformations. d. Coverage of generated conformations with increased sampling on an NVIDIA Titan Xp, 90% confidence interval is shown. e. Coverage of SliM conformations with increased sampling on an NVIDIA Titan Xp, 90% confidence interval is shown. f. Comparison of 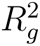 distribution KL divergence. g. Comparison of pairwise C*α* distance distribution KL divergence. h. Comparison of coverage of SliM conformations. i. Comparison of coverage of generated conformations. j. Comparison of average minimum RMSDs of SLiM conformations. k. Comparison of average minimum RMSDs of generated conformations.

As baselines, we ran MD simulations in explicit solvent for 20 ns, and simulations using implicit solvation for 100 ns on the 20 SLiM sequences with the lowest tertiary contacts per residue. A total of 1000 snapshots were taken as the ensemble for both methods. The explicit solvent MD was found to take on average 4.90 times the wall-time of sampling 1000 conformations we PepFlow, and the implicit solvent MD was found to take on average 3.81 times the wall-time of PepFlow sampling (Figre 5d,e). It is important to note that sampling with PepFlow can be further expedited by altering the ODE solver parameters. Increasing the solver error tolerance can lead to a two to five fold increase in sampling speed (Figure S18a). Despite lower sample quality at higher tolerances, due to a higher number of clashes and chain breaks (Figure S18b), the overall structure of generated backbone conformations remains similar (Figure S18c,d). It is thus possible to perform rapid backbone sampling to attain a general picture of conformations of an input sequence.

Relative to PepFlow and explicit solvent MD, implicit solvent simulations were found to perform poorly in generating the SLiM ensembles. We found that implicit solvent simulations tend to explore conformations not represented in the experimental ensembles (Figure 4d), and that generated conformations only cover a small number of the ground-truth conformations (Figure 4e). Despite being trained using an implicit solvent forcefield, PepFlow-generated conformations are much more representative of experimental conformations, due to additional training on real peptide conformations and conformations derived from explicit solvent MD. PepFlow was furthermore found to generate a more representative ensemble than explicit solvent MD, in a much shorter running time (Figure 4d,e). In fact, the coverage of experimental conformations with MD begins to plateau at the 20 ns mark, and the frequency of experimentally represented conformations decreases with increased sampling. Comparatively, the sampled ensemble with PepFlow shows higher concordance in 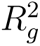 distributions (Figure 4f) and higher concordance in the pairwise C*α* distance distributions (Figure 4g). PepFlow ensembles additionally have higher coverage of experimental conformations (Figure 4h,j) and higher representation of the generated conformations in the experimental ensemble (Figure 4i,k), demonstrating the value of direct sampling of high-likelihood states over trajectory-based methods.

### 2.5 Constrained search in latent space allows modelling of cyclic peptide conformations

Macrocyclization is a technique that is commonly used in therapeutic peptide development as it can confer increased stability, cell membrane permeability and oral bioavailability to linear peptides [25]. While PepFlow is trained to model linear peptides, it is expected that these peptides would assume conformations near the macrocyclic state with some probability. It has previously been shown that exploration of conformations in Boltzmann generator latent space allows for efficient navigation of molecular energy landscapes [13]. On this basis, we implemented a Markov Chain Monte Carlo (MCMC) method in latent space to identify conformations with the target distance between atoms that are covalently bonded in the macrocyclic peptide.

To test this approach, we ran the MCMC on a test set of head-to-tail cyclized peptides from the PDB with 25 starting points each, for 500 iterations. We found that a conformation with the correct distance constraint is consistently generated within 500 iterations (Figure 5a). Comparing the conformations generated with MCMC to 100 conformations sampled directly using PepFlow furthermore shows that the this search effectively moves generation to conformations that more closely resemble the ground-truth (Figure S19a). We additionally compared the performance of the model to AF2 with a positional offset that has recently been shown to allow for the modelling of head-to-tail cyclized peptides [6]. To rank the structures generated by PepFlow, we once again clustered the generated conformations and used the centroid of the most populous cluster as the prediction. On average, the AF2 predictions were more in line with the ground-truth conformations (Figure 5b). We do, however, note that PepFlow consistently generates conformations that are close to the ground-truth (Figure 5c), suggesting that an improved ranking method would significantly increase performance.

**Figure 5:**
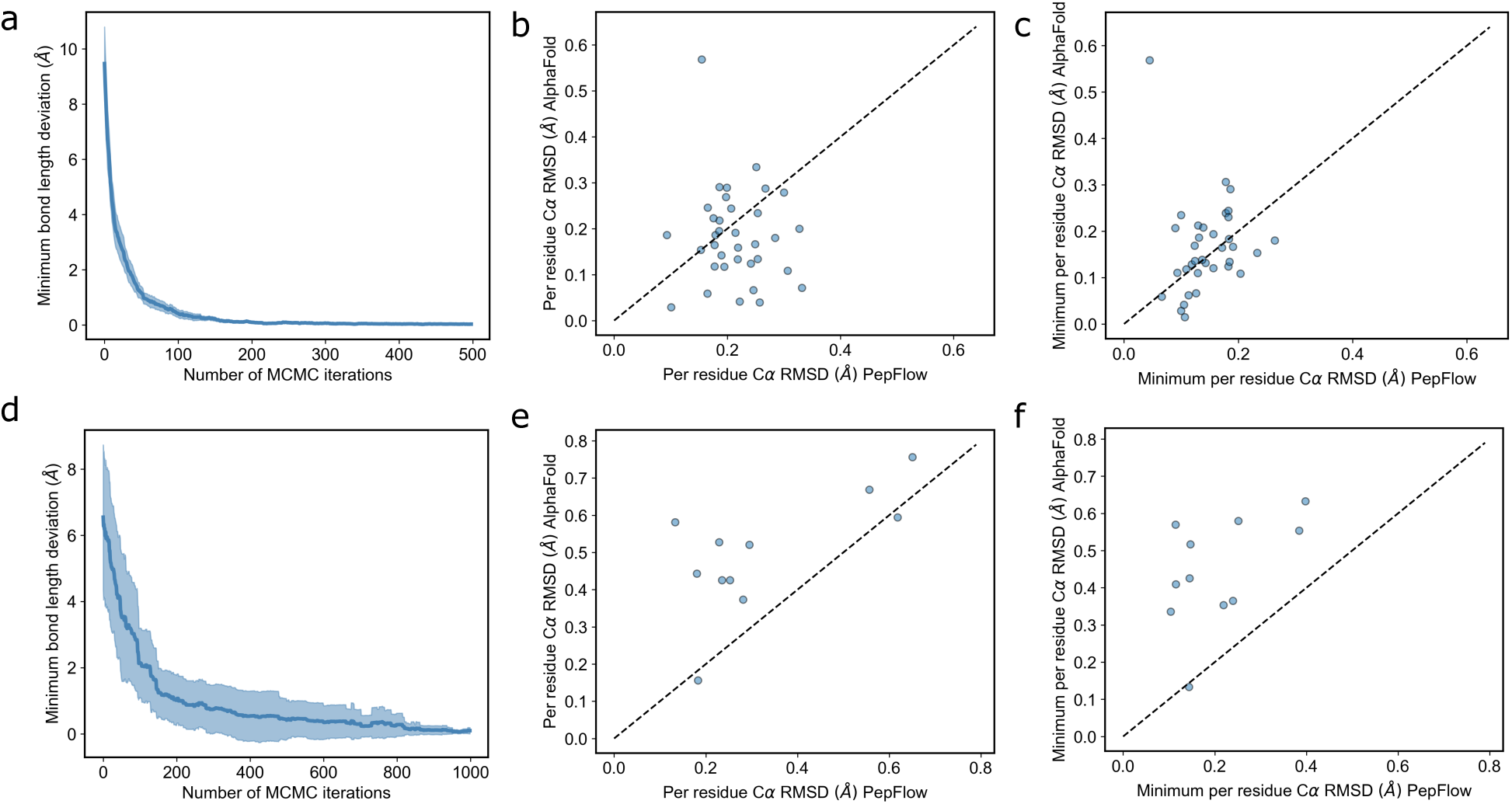
Performance of PepFlow on macrocyclic peptide structure prediction. a. Bond length deviation across MCMC iterations for head-to-tail cyclized peptides. b. RMSD comparison of top ranked AF2 and PepFlow predictions for head-to-tail cyclized peptides. c. Comparison of minimum RMSD of AF2 and PepFlow predictions for head-to-tail cyclized peptides. d. Bond length deviation across MCMC iterations for side chain cyclized peptides. e. RMSD comparison of top ranked AF2 and PepFlow predictions for side chain cyclized peptides. f. Comparison of minimum RMSD of AF2 and PepFlow predictions for side chain cyclized peptides.

One key advantage that PepFlow has compared to AF2 is that side chain atoms are explicitly modelled. Consequently, PepFlow is uniquely able to model peptides with side chain cyclization. We applied the MCMC approach to generate conformations for peptides in the PDB with side chain cyclization. We once again found that the MCMC succeeds in identifying conformations with the desired distance constraint (Figure 5d). We do, however, find that the direct sampling with PepFlow generates conformations with similar backbone structure to the cyclized conformations, suggesting that the target backbone structures are more thermodynamically accessible (Figure S19b,c). Despite this, we find that AF2 fails to accurately capture these conformations, and that the predictions with PepFlow drastically outperform linear peptide prediction with AF2 (Figure 5e,f). The main observed failure case for PepFlow is a set of lasso peptides that assume an unusual fold (Figure S20). The folding of these peptides is catalyzed by specific enzymatic pathways [26] and is thus not directly comparable to the peptides on which PepFlow was trained. PepFlow does accurately predict the structure of the cyclic ring, however, despite generating a more elongated conformation overall (Figure S20).

## 3 Discussion

In this work, we introduced, to the best of our knowledge, the first generalized Boltzmann generator that explicitly models all the degrees of freedom in generated conformations. To mediate faster sampling, this model, PepFlow, makes use of a hypernetwork to generate sequence-specific parameters and contains three networks that progressively model all the atoms in an input peptide sequence. We show that this model is able to effectively generate peptide conformations, and using a simple heuristic to rank conformations allows single-structure prediction that outperforms the state-of-the-art. We furthermore show that PepFlow can efficiently recapitulate experimental ensembles of SLiMs.

In general, the ability to accurately and efficiently sample peptide conformations has the potential to drastically improve peptide docking and design. Peptide docking approaches commonly start with a library of peptide conformations that are docked to a target protein [27, 28]. More accurate generation of peptide ensembles would likely improve this process. PepFlow can furthermore be used to assess the propensity of different sequences to assume conformations that occur on the interface of targeted protein-protein complexes. This could in turn be used to design inhibitory peptides.

We additionally show that latent-space exploration can be used to identify peptide conformations under defined constraints. We use this approach to identify cyclic peptide conformations, but this could, in principle, be to model other constraints such as those in stapled peptides or peptidic fragments between two anchor points. This method could, for example, be used to model the conformations of complementarity-determining regions in antibodies, which have been shown to exhibit peptide-like characteristics [29].

The developed model does, however, contain some limitations that could be improved upon in the future. The current model is trained using an implicit-solvent force field. While PepFlow does accurately capture peptide conformations, performance would likely improve if solvent atoms are explicitly modelled. The modular nature of the PepFlow framework would allow for the incorporation of an additional solvation model. Training the model to enable the direct inclusion of distance constraints would furthermore allow for faster modelling of cyclic structural ensembles, and may improve ranking of generated conformations. Extending the framework to allow for the modelling of the ensemble of peptides bound to a globular protein is an additional step that would also increase the utility of PepFlow. Overall, the developed framework is highly flexible and serves as a valuable proof-of-concept for all-atom modelling through deep learning.

## 4 Methods

### 4.1 Score-based generative framework

We initially train our models in a continuous-time diffusion framework following Song et. al 2021 [16]. Input data *x* is noised to a prior distribution according to an SDE with the following form:

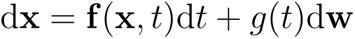

Where *t* ∈ [0, 1] denotes the time, **f** (*., t*) is the drift coefficient, *g*(*t*) is the diffusion coefficient and d**w** is the standard Wiener process. Given the forward diffusion process, the following reverse-time SDE can be used to denoise the data:

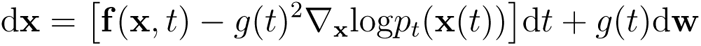

The learn this reverse process, a network, **s***_θ_*(**x***, t*) is trained to estimate the score.

PepFlow includes three such score networks. The first network, *s_θ_s__* (*., t*) learns to generate coordinates of the backbone atoms and side chain centroids, **x***_b_* and **x***_c_*, respectively, by denoising relative atomic coordinates. Here, a subset of the parameters in the network are predicted by a hypernetwork based on an input peptide sequence. The second score network, *s_θ_r__* (*., t|***x***_b_*(0), **x***_c_*(0)), denoises the coordinates of side chain heavy atoms, **x***_r_* based on the backbone atoms and centroids, while enforcing that the side chain centroid is maintained. The final network, *s_θh_* (*., t|***x***_b_*(0), **x***_r_*(0)), denoises the relative coordinates of hydrogen atoms to their bound heavy atom to allow for peptide protonation.

The backbone and protonation noising processes make use of the sub-Variance Preserving (sub-VP) SDE defined in Song et al. 2021 [16]:

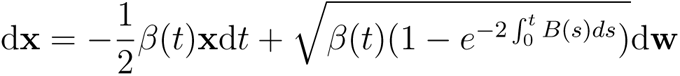

Where *B*(*t*) represents a linear interpolation between two hyperparameters (Table S1). We make use of a CoM preserving variant of the sub-VP SDE to define the noising process for the rotamer model:

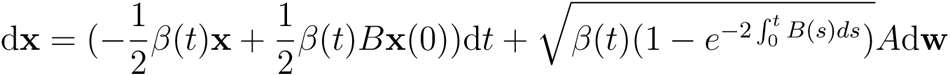

Here, the matices *A* and *B* preserve the centroid positions for each side chain (Supplementary Information).

### 4.2 ODE probabality flow

One of the defining features of SGMs is the existence of a probability flow ODE that represents a deterministic process with same probability densities across its trajectory [16]:

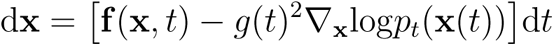

Defining a flow with an ODE addtionally allows exact-likelihood computation using the instantaneous change instantaneous change of variables formula [30]. With **f***_θ_*(**x***, t*) = **f** (**x***, t*)*− g*(*t*)^2^∇**_x_**log*p_t_*(**x**(*t*)):

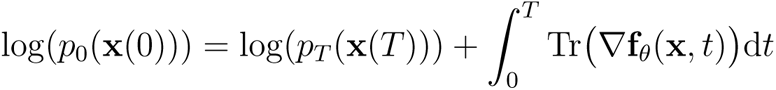

Where Tr(∇**f***_θ_*(**x***, t*) denotes the trace of the Jacobian of **f***_θ_*(**x***, t*). Computing the exact trace is computationally expensive. We thus use the Hutchinson’s estimator of the trace as previously described, which can be computed efficiently as a vector-Jacobian product [30]:

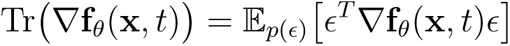

Where ℝ[*ɛ*] = 0 and Cov(*ɛ*). Here, we use a standard Rademacher distribution to sample *ɛ*. *p_T_* (**x***_b_*(*T*), **x***_c_*(*T*)) follows a standard 3(N-1)-dimensional Gaussian distribution, where N is the number of backbone atoms and the number of centroids, accounting for the invariance of the likelihood to the position of the N-terminal nitrogen. *p_T_* (**x***_r_*(*T*)) is modelled as a centered Gaussian distribution (Supplementary Information), and *p_T_* (**x***_h_*(*T*)) follows a standard Gaussian. Using this, the marginal log-likelihood across all atoms can be computed:

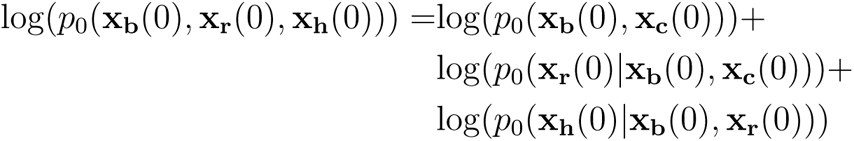

This holds due to the fact that log(*p*_0_(**x_b_**(0), **x_c_**(0), **x_r_**(0))) = log(*p*_0_(**x_b_**(0), **x_r_**(0))), as the side chain centroids are preserved in the rotamer generation flow.

### 4.3 Hypernetwork architecture

The hypernetwork takes as input a peptide sequence and generates sequence-specific parameters for the main network. A token is concatenated to the provided sequence that stores a high dimensional representation of the input peptide, as done in classification tasks with large language models [31]. The resulting sequence goes through multiple attention blocks consisting of layers of standard scaled dot-product self-attention [32]:

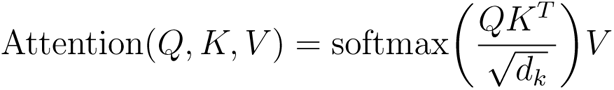

Where *Q, K, V* are linear projections of the sequence and *d_k_* is the size of the embedding dimension of *Q* and *K*. The outputs of the self-attention layers are then passed through feed-forward networks [32]. In total, the hypernetwork consists of three attention blocks, with the embedding dimension of *Q, K*, and *V* set to 128, the hidden dimension of the feed-forward networks set to 64, and the dropout rate set to 0.3.

For every linear layer in the main-network, a feed-forward network projects the generated peptide embedding to a weight matrix and bias vector. The feed-forward networks consist of an initial projection layer followed by a ReLU, and then projection to the output matrices. The previously described hyper fan-in method is used to initialize the layers [33].

### 4.4 Main-network architecture

For the dynamics networks of PepFlow, we adapt EGNN layers [17, 34]. Each layer consists of a series of feed-forward networks with softplus non-linearities that operate on atom features **a**, edge features **e**, coordinates **x** and time *t*. For a layer *l*,

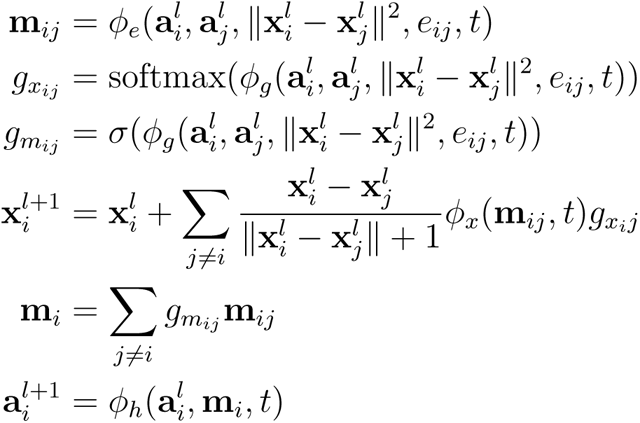

Each feed-forward network in the model conditions on *t* using an affine transformation. More specifically, for any particular layer we compute the output as *ϕ*(*., t*) = *σ*(Linear(*t*))FFN(.) + tanh(Linear(*t*)). The exact model hyperparameters are listed in Table S1.

In the case of the backbone model, a fully-connected graph was used. In the case of the rotamer model, each residue had edges to other residues with atoms that were within 8 Å of the side chain centroid. The rotamer model additionally centers the side chain atoms after each layer to maintain the predicted centroid. Finally, edges in the graph used by the protonation model were present between hydrogen atoms and other hydrogen atoms bound to the same heavy atom, as well as the heavy atoms that were within 4 Å of the bound heavy atom.

### 4.5 Input features

Each atom has three input features, the sequence position of the input amino acid, the identity of the input amino acid, and the atom identity. Each of these features are represented as one-hot encodings. All hydrogen atoms are given the same atom identity encoding, and atom pairs that are found in identical chemical environments are given the same encoding, to enforce permutation invariance (Table S2). In total, the atom identity one-hot encoding includes 43 classes. In the graph edges, the presence or absence of a covalent bond connecting two atoms is included as a binary feature.

### 4.6 Training by example

The score networks are trained to minimize the following weighted denoising score matching (DSM) loss:

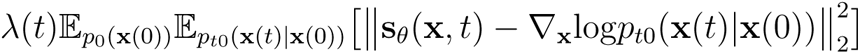

Where 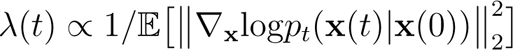. An Adam optimizer is used during training with *β*_1_ = 0.9*, β*_2_ = 0.999 and *ɛ* = 1e 8. To stabilize training, an exponential moving average (EMA) was applied to the weights, as done previously [35], and the norm of the gradients on the parameters is clipped. Full training parameters are listed in Table S3.

### 4.7 Training by energy

During training by energy, the three score networks are jointly optimized to minimize the KL divergence between the PepFlow proposal distribution and the Boltzmann distribution paramaterized by a molecular forcefield. Following the derivation of Nóe et al. (2019) [13], with *q_T_* being the probability distribution of the Boltzmann distribution, paramaterized by the energy function *u*(.), and transformed with the reverse-time ODE flow:

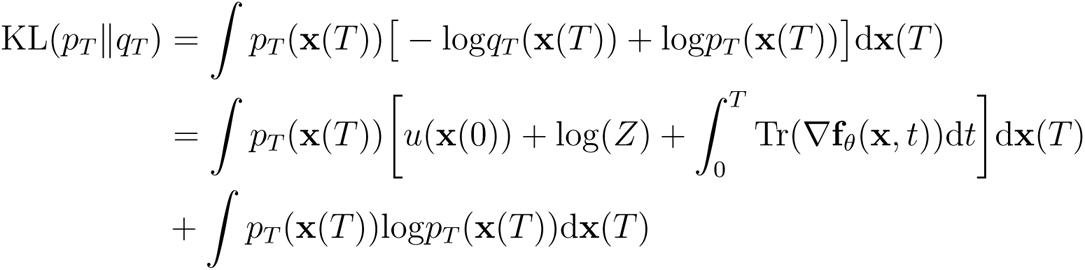

Where *Z* is the partition function for the Boltzmann distribution. Removing the terms that do not depend on the model parameters yields the following loss function:

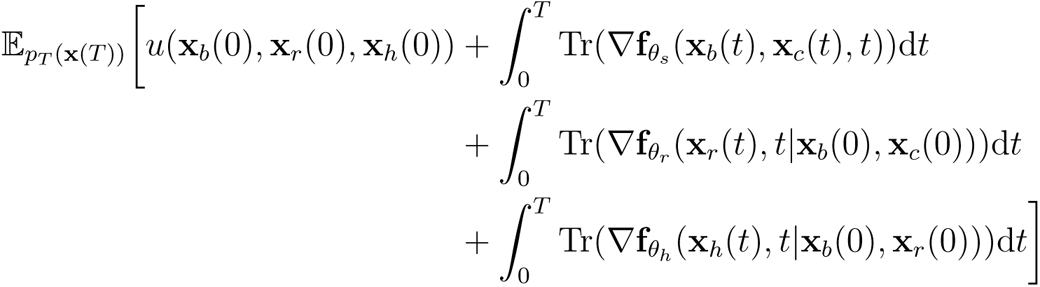

At every training iteration, eight conformations for an input peptide are sampled. Sampling and likelihood computation are done using the RK23 solver from `scipy.integrate.solve_ivp` [36], with atol=1e-3 and rtol=1e-3, wrapped using the adjoint solver from the `torchdiffeq` package [30]. To solve the reverse-time ODE during backpropogation with the adjoint method, the fixed-step rk4 solver from the `torchdiffeq` package is used with a step size of 5e-2. Energies are evaluated using the Amber ff99SB-ILDN protein force field [37], with the GB-Neck2 implicit solvent model [38] using the `openmm` package [39]. A learning rate of 1e-5 is used, with a batch size of 1 and the gradient norm clipped at 100 on the KL loss. An additional DSM loss is added where 16 conformations are sampled from a set of MD simulations at each iteration. Training for 1000 iterations on a single NVIDIA A100 took approximately three days.

### 4.8 MD simulations

Linear peptides generated using chimera were used as starting conformations for MD simulations [40]. The resulting conformations were then relaxed in `openmm`, before running the production step with a step size of 2 femtoseconds and a temperature of 310 K. A Langevin integrator is used with a friction coefficient of 0.1 picoseconds. Implicit solvent MDs were run with the Amber ff99SB-ILDN protein force field [37] and the GB-Neck2 implicit solvent model [38]. For explicit solvent simulations, the peptides are solvated in TIP3P water molecules at a padding of 12 Å, as done in DBAASP simulations [19].

### 4.9 Conformational sampling

Conformational sampling is done iteratively, starting with the backbone and centroid coordinates, followed by sampling the side chain heavy atom coordinates and finally peptide protonation. Sampling is done with the RK23 solver from scipy.integrate.solve ivp [36] using atol=1e-4 and rtol=1e-4. For faster sampling, atol and rtol are set to 1e-2 for back-bone generation and 1e-3 for side chain and proton generation. For likelihood computation, parameters of atol=1e-3 and rtol=1e-3 are used across all three steps.

Given that PepFlow modules are invariant to reflection, the generated conformations occasionally assume D-amino acid conformations instead of L-amino acid conformations. The model does overwhelmingly generate conformations that have consistent chirality throughout (Figure S21). Chirality of each conformation is estimated by whether the following value is less than 0:

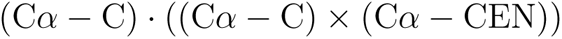

Where C*α* is the coordinates of the C*α* atom, C is the coordinates of the C atom and CEN is the coordinates of the side chain centroid. Conformations with less than 50% L-amino acids are reflected along the x-axis.

When doing single-structure prediction, 100 conformations are sampled. Hierarchical clustering is then done on the generated conformations using the average linkage method from `scipy.cluster.hierarchy.linkage` [36], using pairwise C*α* RMSDs computed using `mdtraj` [41]. Clusters are then separated at at a distance threshold of 1.5 Å, and the centroid of the most populated cluster is used as the predicted conformation.

### 4.10 Sampling cyclic peptide conformations

To sample cyclic peptide conformations MCMC searches were performed in model latent space. The Metropolis algorithm was used with the objective of minimizing the deviation in the ideal bond length of the cyclic bond and the distance between the two atoms, **x***_i_* and **x***_j_*, forming that bond in the sampled conformations. To avoid collapsing the generated conformations, a term was included to penalize clashes. This resulted in the following Metropolis criterion:

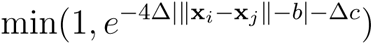

Where *b* is the ideal bond length and *c* is the number of clashes, defined as the number of non-covalently-bonded atoms within 1.6Å of each other. At each step, **x**(*T*) is perturbed with Gaussian noise and a step size of 0.2. In total, 500 steps were performed for head-to-tail cyclization and 1000 steps were performed for side chain cyclization, with 25 starting points. We perform initial steps with atol and rtol set to 1e-2 in the ODE solver, based on the fact that despite the lower sample quality, the overall structure of generated conformations are similar (Figure S18). Every 25 iterations, we assess whether decreasing the error tolerance changes the bond length by more than 0.25 Å, and if so, the error tolerance is decreased. The generated conformations with less than a 1.5 Å deviation from the ideal bond length are clustered and the centroid of the most populous cluster is used as the final prediction. The clustering threshold is set to 1.5 Å and increased incrementally by 0.1 Å until a cluster is generated with more than one sample.

### 4.11 PDB datasets

To generate training datasets from the PDB, all structures with accessions earlier than 14/09/2021 were downloaded. The protein fragment dataset was generated from crystal structures lacking a protein chain of less than 16 amino acids. PDB entries were filtered by resolution at a threshold of 3 Å and the remaining protein chains were clustered using MMSeqs2 at 30% identity [42]. The centroids of the resulting clusters were fragmented at lengths 3 to 15, resulting in approximately 40 million unique fragment sequences. 90% of the fragment sequences were randomly selected for training, and of the remaining fragments in the validation set, 1000 were selected per sequence length for model comparisions and hyperparameter tuning. To account for the comparatively low amount of strand-forming fragments, sequences were weighted by their secondary structure to sample an equal number of helices, strands and coils while training the backbone model.

The peptide conformation dataset additionally includes NMR structures. Structures with non-natural amino acids were filtered out. Cyclic peptides were separated from linear peptides based on the presence of a non-disulfide covalent bond between non-adjacent residues. For training, linear peptides with accession dates earlier than 14/09/2021 were clustered at 30% identity and 10% sequence coverage using MMSeqs2 [42]. 10% of the clusters were used as a validation set, with the remaining was used as the training set. For testing, sequences of peptides from recently added PDB entries were used. Sequences that were redundant with either the peptide training and validation sets or the fragment training and validation sets were filtered out. This was done at a 30% sequence identity threshold and 10% sequence coverage threshold based on MMseqs2 search [42]. The result was a test dataset of 167 linear peptides, and a test dataset of 46 cyclic peptides.

### 4.12 MD dataset

MD simulations of peptides with lengths 8 to 15 were downloaded from the DBAASP dataset [19]. Sequences were filtered by redundancy with the sequences in the training and validation sets at a 30% identity and 10% sequence coverage threshold. To avoid high redundancy within the MD dataset itself, sequences were clustered at 30% identity and 10% sequence coverage, and non-centroid sequences were removed. 80 sequences that were redundant with the training sequences were further split and used as training and validation datasets for initial training the protonation model, and a validation dataset for the backbone model. 50% of the remaining 802 simulations lacking C-terminal or N-terminal modifications were used a training dataset in the training by energy phase, and the remainder were used as a test dataset for benchmarking.

### 4.13 SLiM dataset

SLiMs in protein sequences from the PED [23] were identified using the ELM server [24]. Identified SLiMs from lengths 8 to 15, with at least 50 models, were filtered for redundancy with the training and validation datasets and clustered. The 572 remaining sequences were extracted from their respective PED models. To identify tertiary contacts in the extracted fragments, we annotated the number of residues with a heavy atom within 8 Å of the fragment residues that was more than three residues away in primary sequence and was not within the fragment. After filtering by fragments with fewer than three tertiary contacts per residue, 75 sequences remained.

### 4.14 Baseline approaches

We ran AF2 with full MSA generation, using default settings, and no templates. To generate head-to-tail cyclized conformations with AF2, we added an offset to the positional encodings as previously described [6]. ESMFold and MODPEP were run with default settings [20, 22]. PSIPRED was used to generate secondary structure predictions for input to MODPEP [43].

## 6 Code availability

The code to run PepFlow is available at https://gitlab.com/oabdin/pepflow.

## 8 Supplementary Information

### 8.1 Denoising relative atomic coordinates

Backbone coordinates are denoised as relative coordinates. More specifically, every atomic coordinate is encoded relative to its N-terminal neighbor. Centroid atom positions are encoded relative to the C*α* of the corresponding residue. This noising process is equivalent, up to a scaling constant, to a previously described ideal chain covariance model [1]. Using the previously defined sub-VP SDE [2] for the noising process, and with absolute coordinates **x**, we can define the distribution of the relative position between atoms *i*^th^ and *j*^th^ in the backbone as follows:

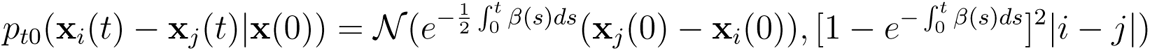

With 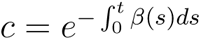, the expectation of squared distances is as follows:

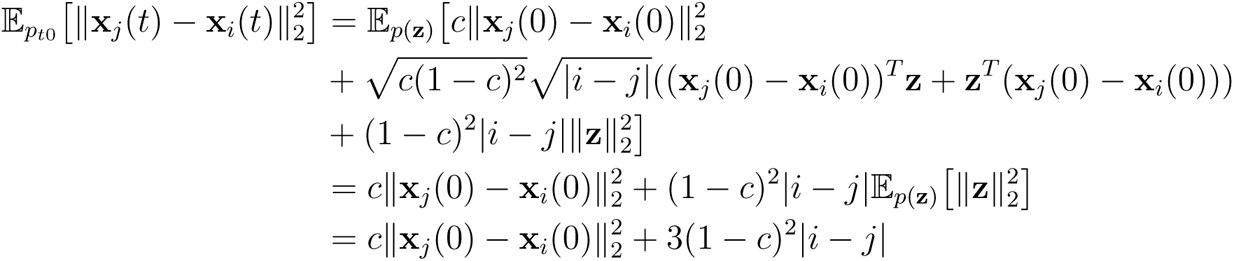

This expectation is of a similar form to the expression derived in Ingraham et al. 2022 [1], and leads to the following expectation for the radius of gyration, 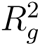, at time *T* :

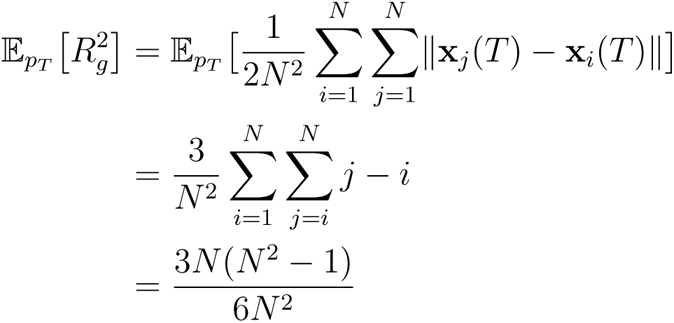

For large values of *N*, this follows ideal-chain scaling.

### 8.2 Center-of-mass perserving SDE

To develop an SDE that preserves sampled centroids, we extend previous analysis on equivariant normalizing flows [3, 4]. Let *x* define a system with a dimensionality of *d*, and *k* sets of coordinates that each have a set CoM so that

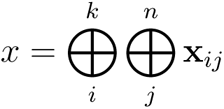

Where ⊕ denotes concatenation. For a vector **x***_ij_* ∈ ℝ*^N_i_^*, the following matrix removes the mean:

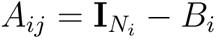

Where

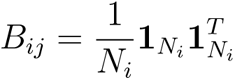

Then the following block-diagonal matrix that removes the CoM for each set of coordinates:

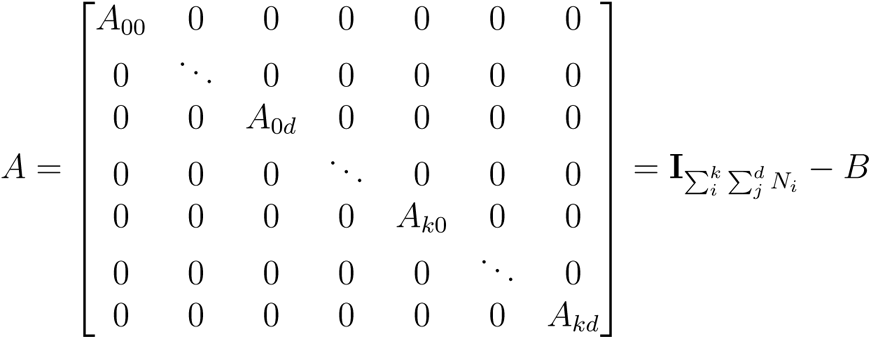

The following is a CoM preserving variant of the sub-VP SDE [2]:

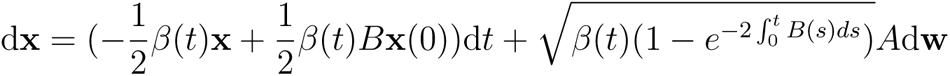

Given the affine drift and diffusion coefficients, it is possible to compute a closed-form for the probability distribution *p*_0_*_t_*(**x**(*t*) **x**(0)) using the differential equations defined in Säarkkä and Solin (2019) [5]:

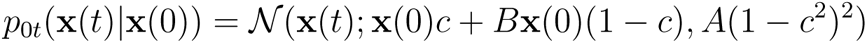

Where 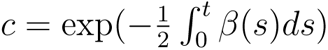. Given that *A*^2^ = *A* and *A^T^* = *A* this is equivalent to

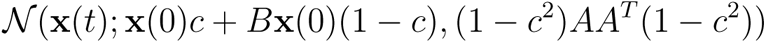

Therefore sampling from this distribution can be done by projecting sampled noise with *A*. Importantly, for *x*(*t*) sampled from this distribution

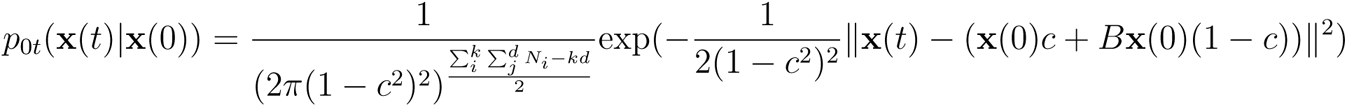

The closed form solution of *p*_0_*_t_*(**x**(*t*) **x**(0)) allows the exact computation of **_x_**_(_*_t_*_)_log(**x**(*t*) **x**(0)) during training.

### 8.3 Supplementary Tables

**Supplementary Table 1:** Hyperparameters of PepFlow dynamics networks

| Model | Number of predicted layers | Number of optimized layers | Hidden dimension | SDE | $\beta_{\min}$ | $\beta_{\max}$ |
| --- | --- | --- | --- | --- | --- | --- |
| Backbone | 4 | 4 | 256 | sub-VP SDE | 0.1 | 20 |
| Rotamer | 0 | 4 | 128 | CoMP sub-VP SDE | 0.1 | 20 |
| Protonation | 0 | 4 | 64 | sub-VP SDE | 0.01 | 20 |

**Supplementary Table 2:** Ambiguous atoms

| Amino acid three-letter code | Atom one | Atom two |
| --- | --- | --- |
| Asp | OD1 | OD2 |
| Glu | OE1 | OE2 |
| Phe | CD1 | CD2 |
| Phe | CE1 | CE2 |
| Tyr | CD1 | CD2 |
| Tyr | CE1 | CE2 |
| Leu | CD1 | CD2 |
| Val | CG1 | CG2 |
| Arg | NH1 | NH2 |
| - | O | OXT |

**Supplementary Table 3:** Hyperparameters used during training of PepFlow networks

| Model | Learning rate | Batch size | EMA decay | Gradient clip | Iterations (M) | GPU | Walltime (Days) |
| --- | --- | --- | --- | --- | --- | --- | --- |
| Backbone | 1e-4 | 32 | 0.9999 | 1 | 2.12 | NVIDIA A100 | 23.8 |
| Rotamer | 5e-4 | 8 | 0.9999 | 1 | 3.68 | NVIDIA V100<br>Volta | 28.4 |
| Protonation | 5e-4 | 4 | 0.9999 | None | 0.53 | NVIDIA V100<br>Volta | 8.5 |

### 8.4 Supplementary Figures

**Supplementary Figure 1:**
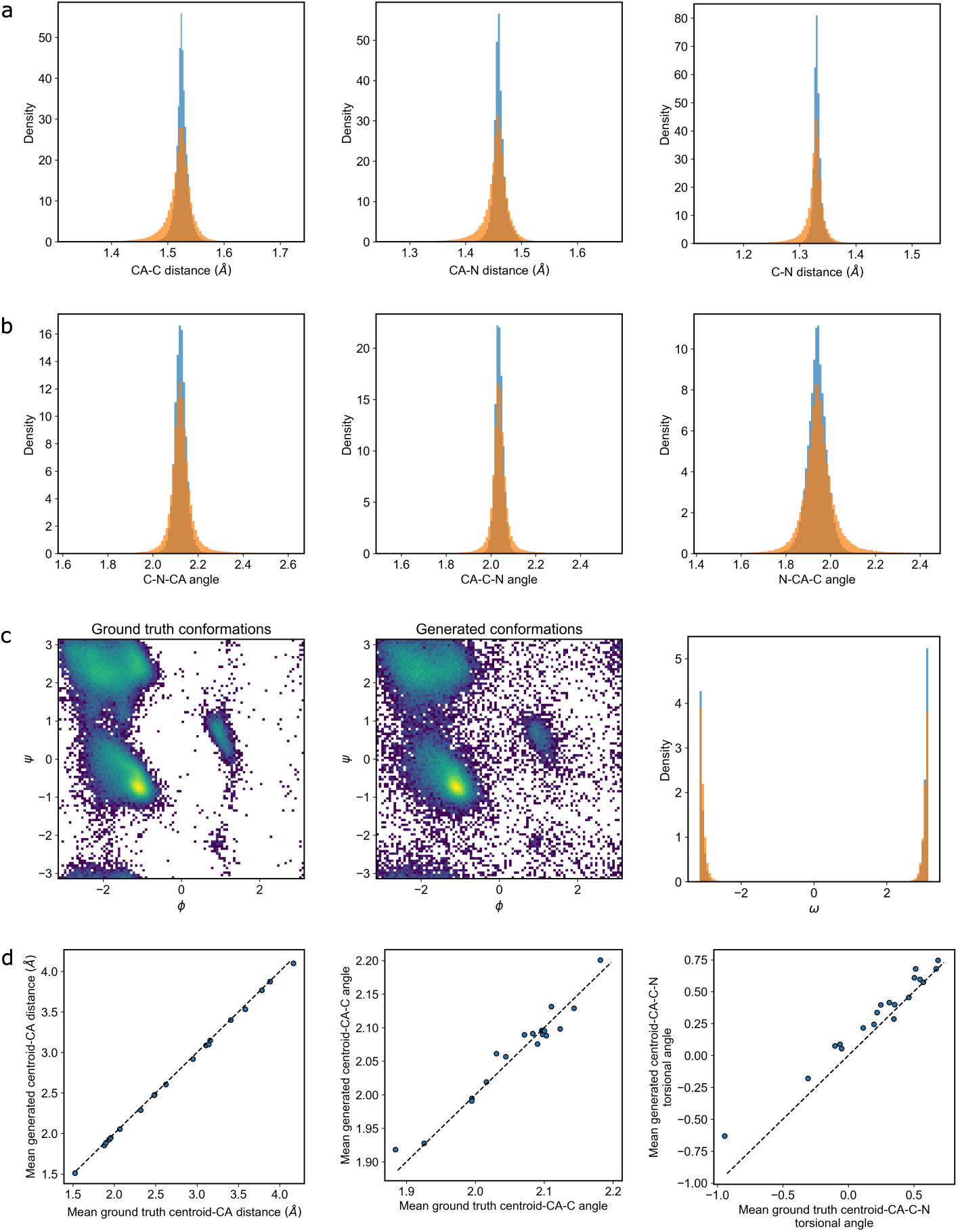
Molecular features of generated peptide conformations. Distributions from ground truth molecules are shown in blue and distributions from generated conformations are shown in orange. a. Backbone bond length distributions. b. Backbone bond angle distributions. c. Backbone torsion distributions. d. Comparison of mean centroid positions across different amino acids.

**Supplementary Figure 2:**
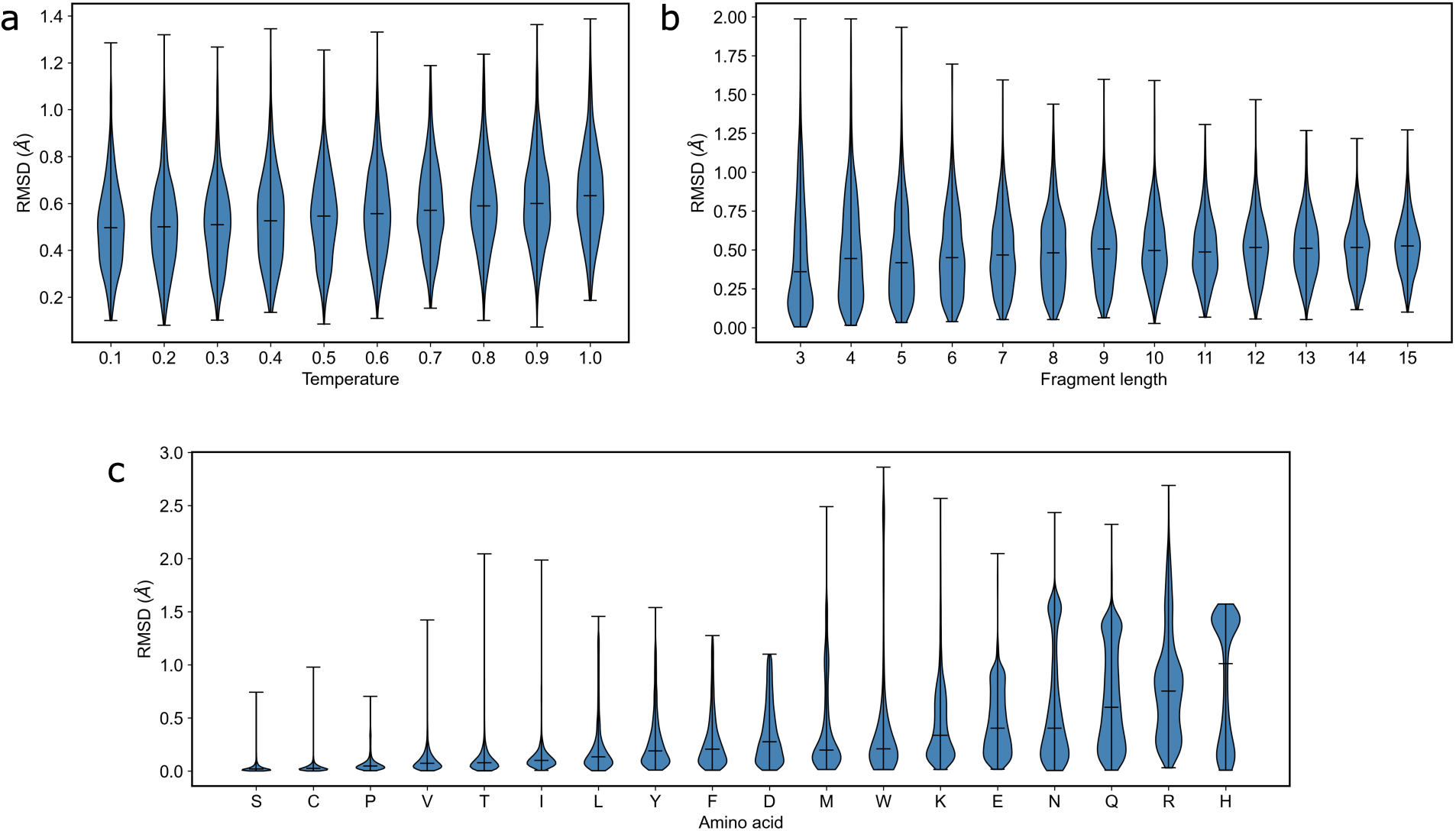
RMSDs of generated side chain conformations to ground truth fragments. a. side chain RMSDs of fragments with length 15 at different with latent temperature scaling. b. side chain RMSDs of fragments of different lengths at a temperature of 0.4. c. side chain RMSDs for different amino acids at a temperature of 0.4.

**Supplementary Figure 3:**
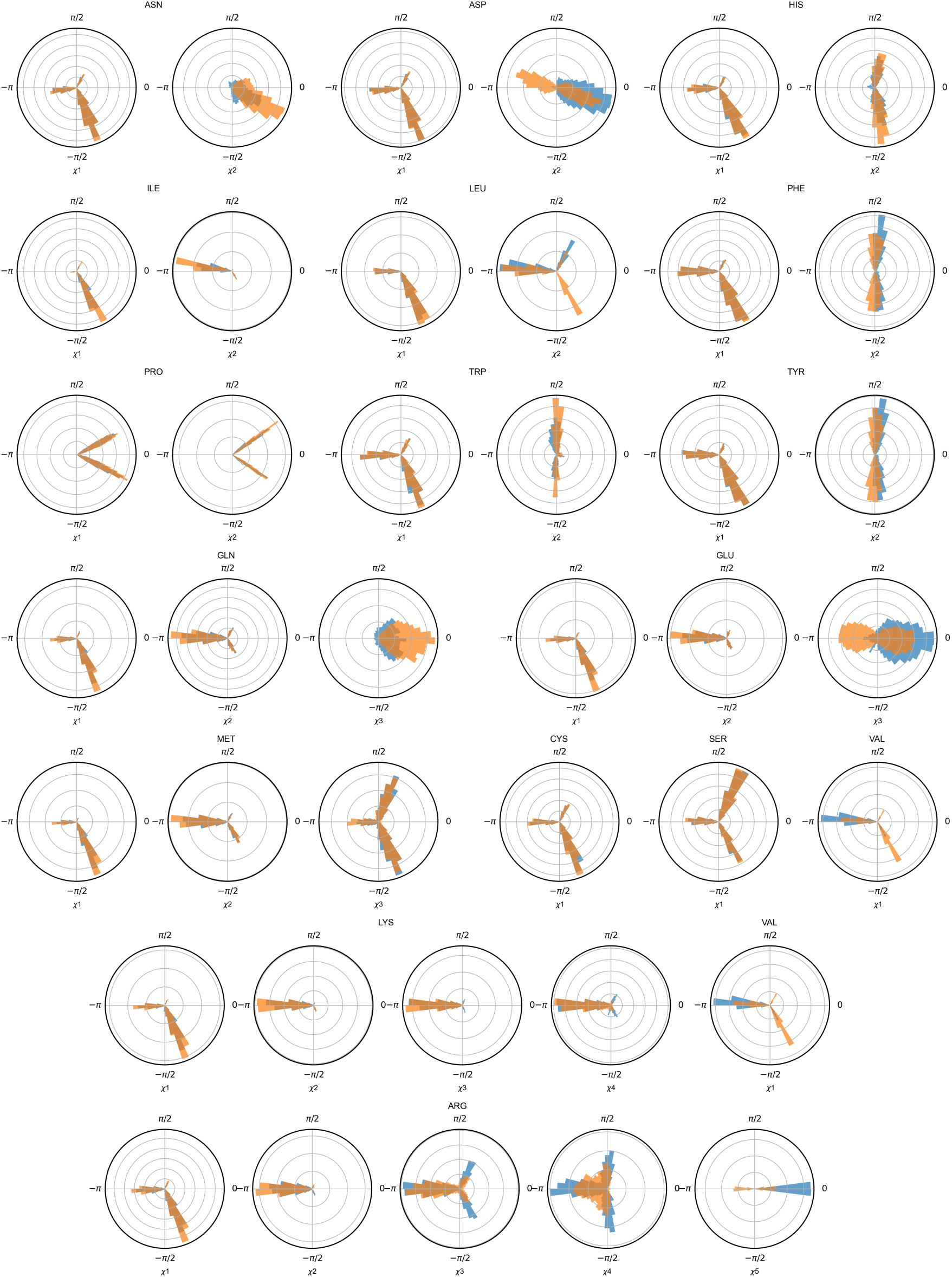
side chain torsion angle distributions of generated peptides. Distributions from ground truth molecules are shown in blue and distributions from generated conformations are shown in orange.

**Supplementary Figure 4:**
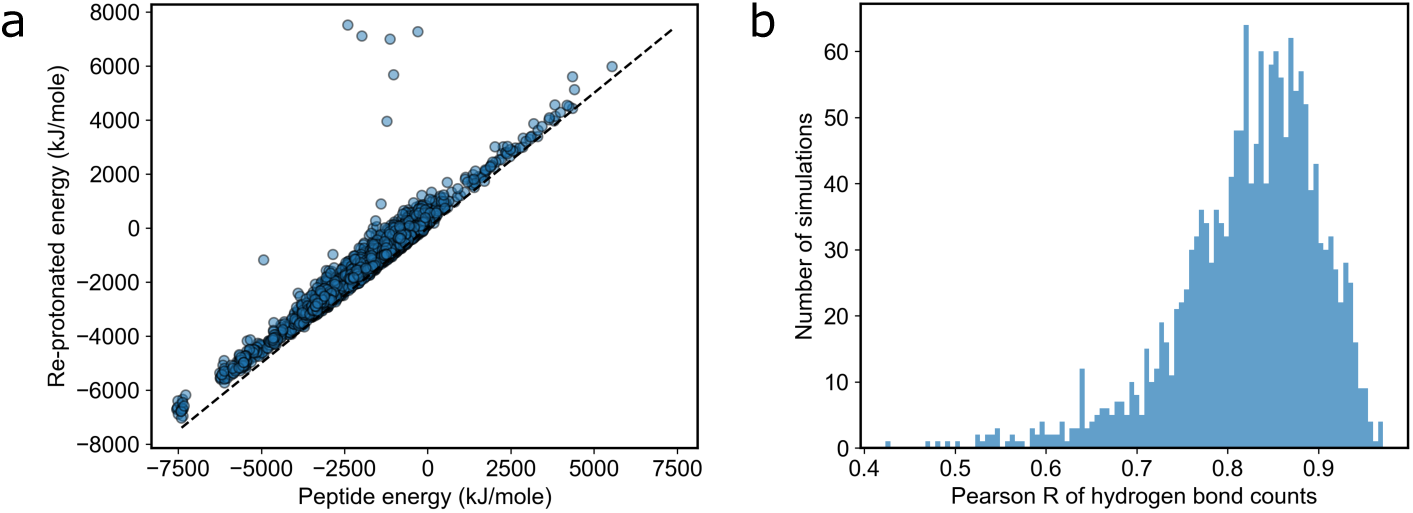
Comparison of conformations from MD simulations with re-protonated conformations. a. Comparison of the estimated energy of conformations from MD simulations with re-protonated conformations. b. Pearson correlations between hydrogen bond counts in MD simulations and re-protonated conformations.

**Supplementary Figure 5:**
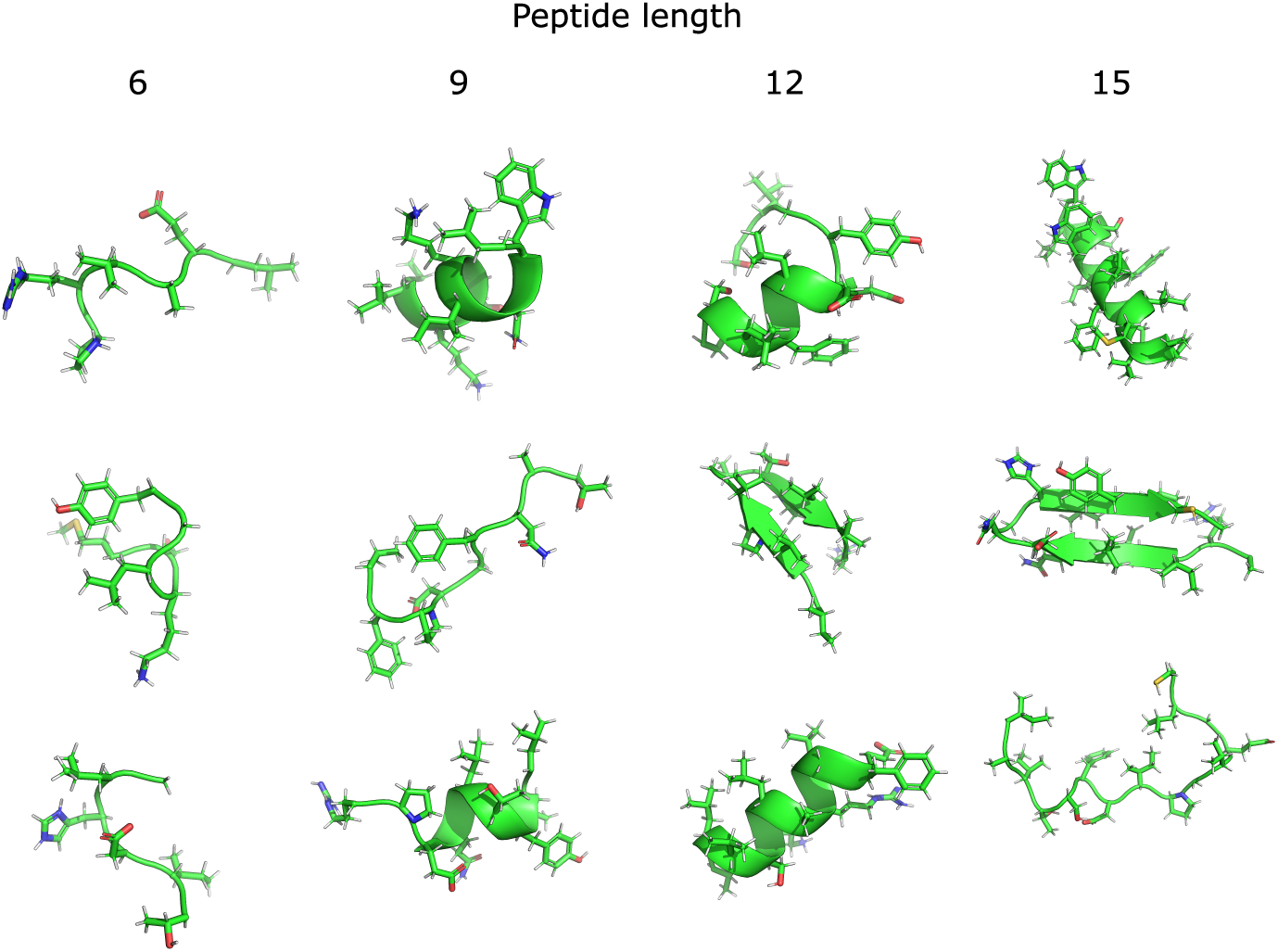
Example PepFlow conformations generated for sequences of different lengths.

**Supplementary Figure 6:**
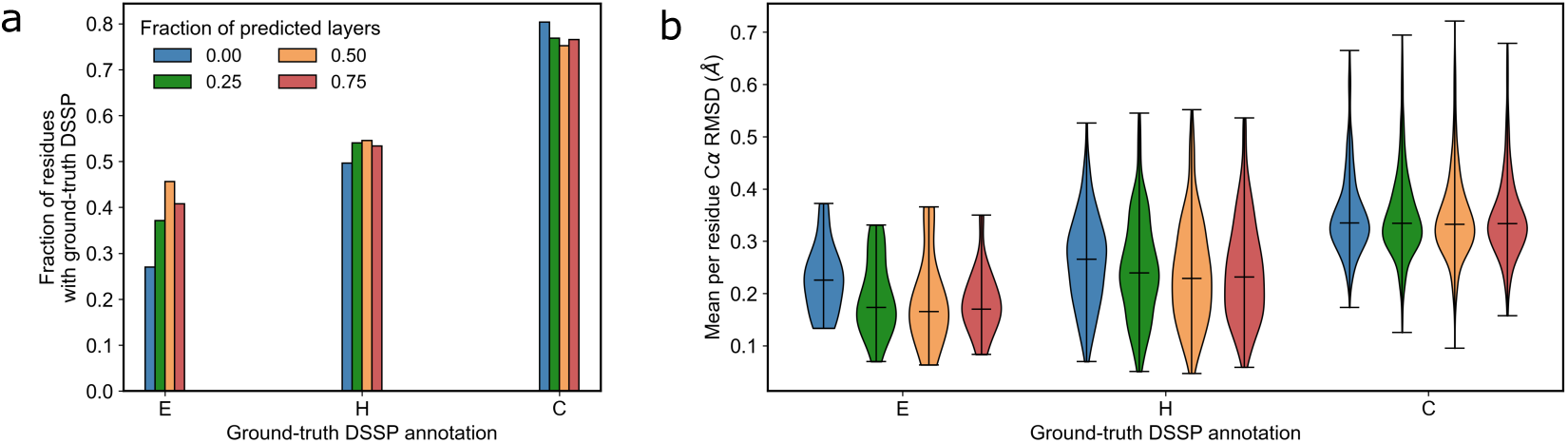
Prediction of fragment structures with varying hypernetwork layers. a. Percentage of residues sampled with the same secondary structure as 15 amino acid fragments from the PDB. b. RMSDs of sampled conformations to 15 amino acid fragments from the PDB.

**Supplementary Figure 7:**
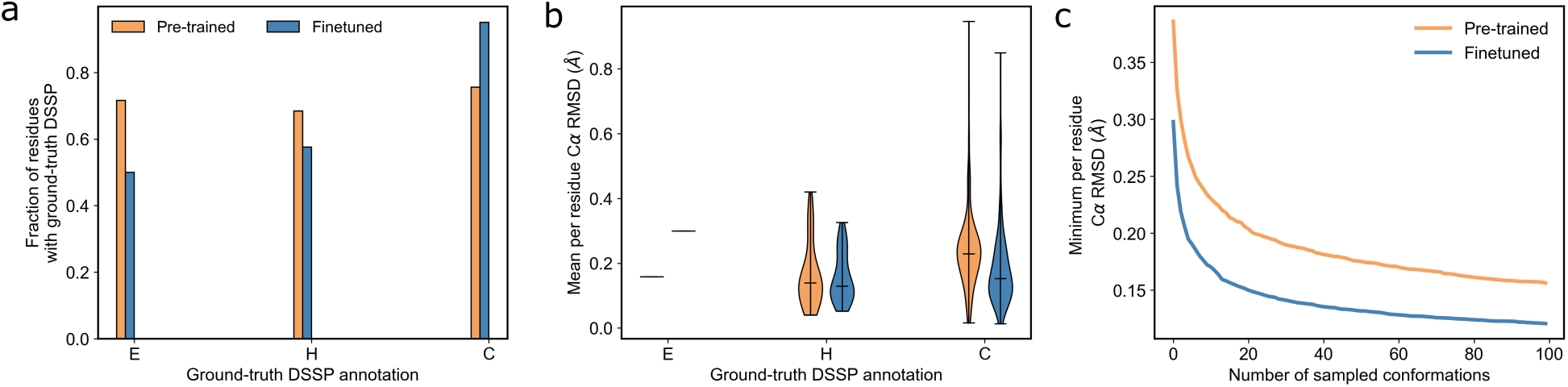
Comparison of pre-trained and finetuned PepFlow on peptide validation set. a. Comparison of secondary structure prediction. b. Comparison of C*α* RMSDs to ground-truth peptide conformations. c. Comparison of RMSDs as more samples are generated with each model.

**Supplementary Figure 8:**
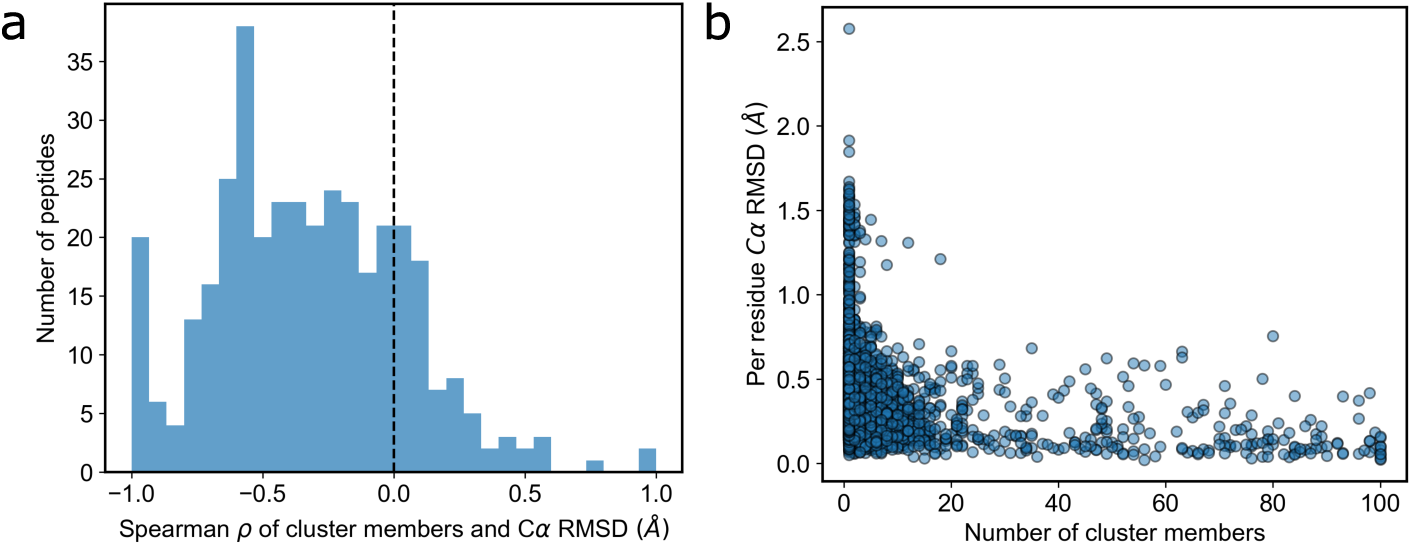
Correlation between number of cluster members and C*α* RMSDs of the cluster centroid. a. Spearman *ρ* for each every peptide in the validation set. b. Overall relationship between number of cluster members and C*α* RMSDs of the cluster centroid.

**Supplementary Figure 9:**
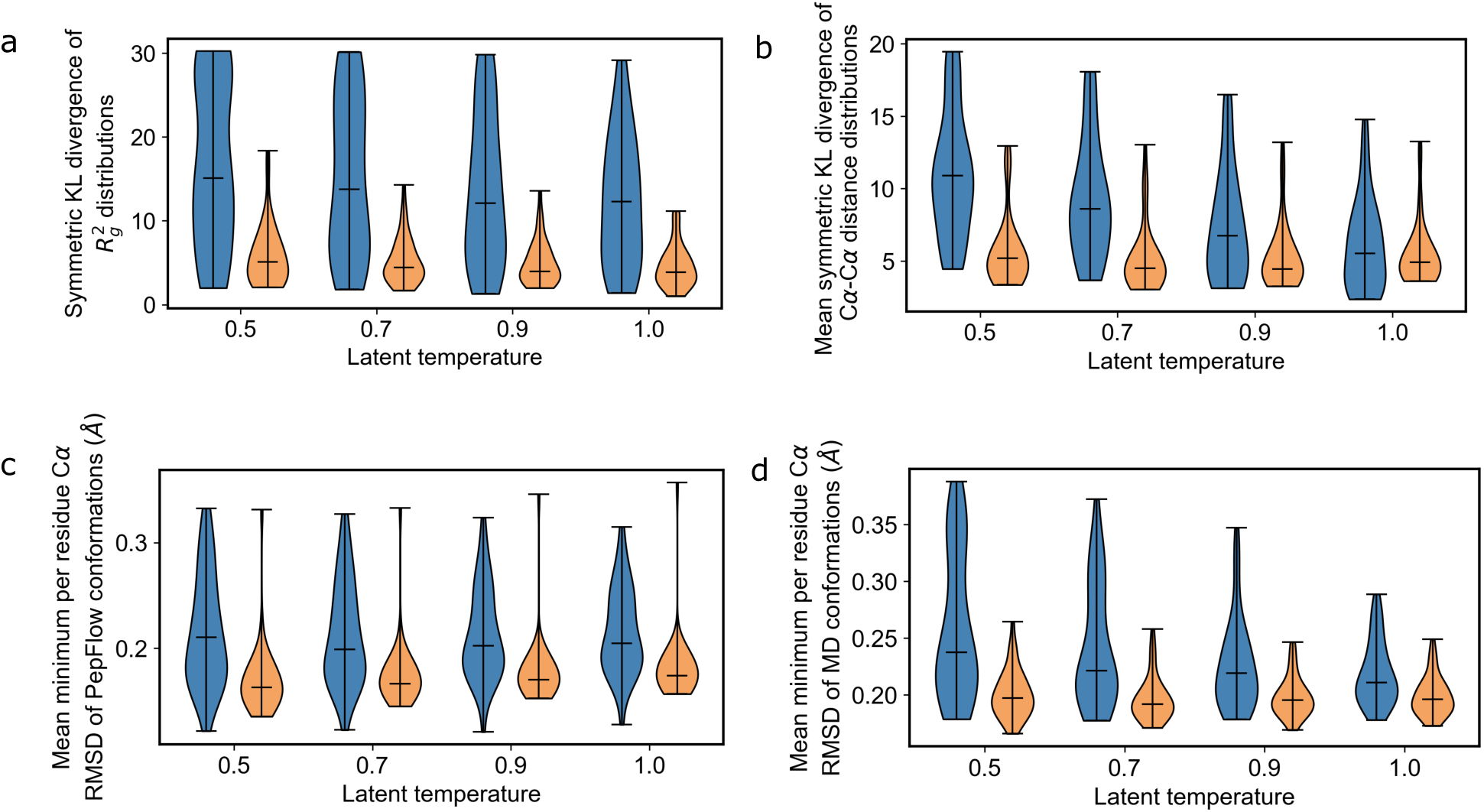
Comparison of performance on ensemble generation before and after training by energy. Results before training by energy are shown in blue. Results after training by energy are shown in orange. a. KL divergence of distributions of radius of gyration between PepFlow generated conformations and conformations in the MD validation set. b. Mean KL divergence of distributions of pairwise C*α*-C*α* distances between PepFlow generated conformations and conformations in the MD validation set. c. Distributions of mean minimum RMSD between PepFlow generated conformations and MD conformations. d. Distributions of mean minimum RMSD between MD conformations and PepFlow generated conformations.

**Supplementary Figure 10:**
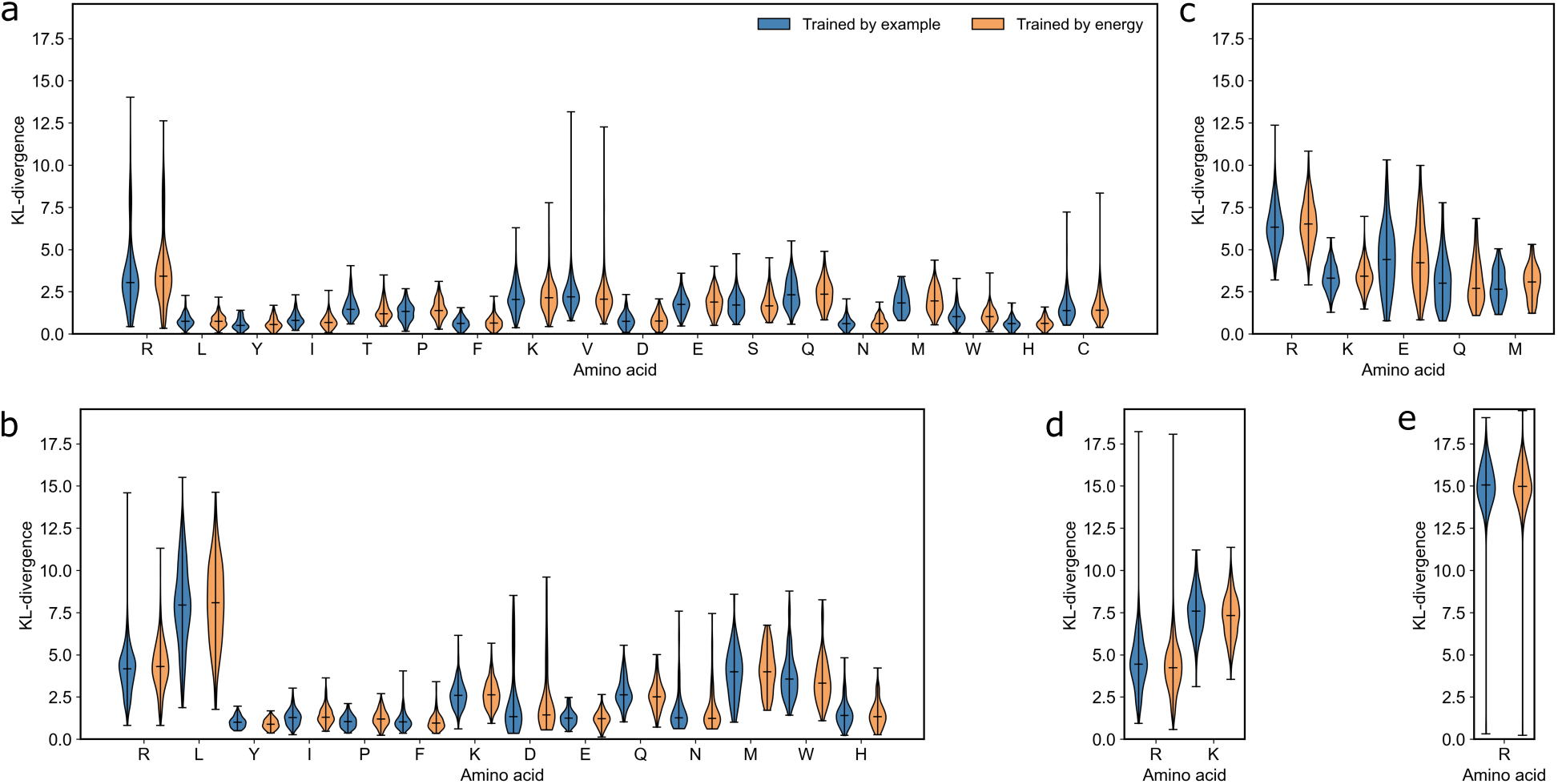
Comparison of rotamer generation before and after training by energy on the MD test set. Results before training by energy are shown in blue. Results after training by energy are shown in orange. a-e. KL divergence between distributions of chi angles 1-5 for generated conformations and conformations from MD simulations across different amino acids.

**Supplementary Figure 11:**
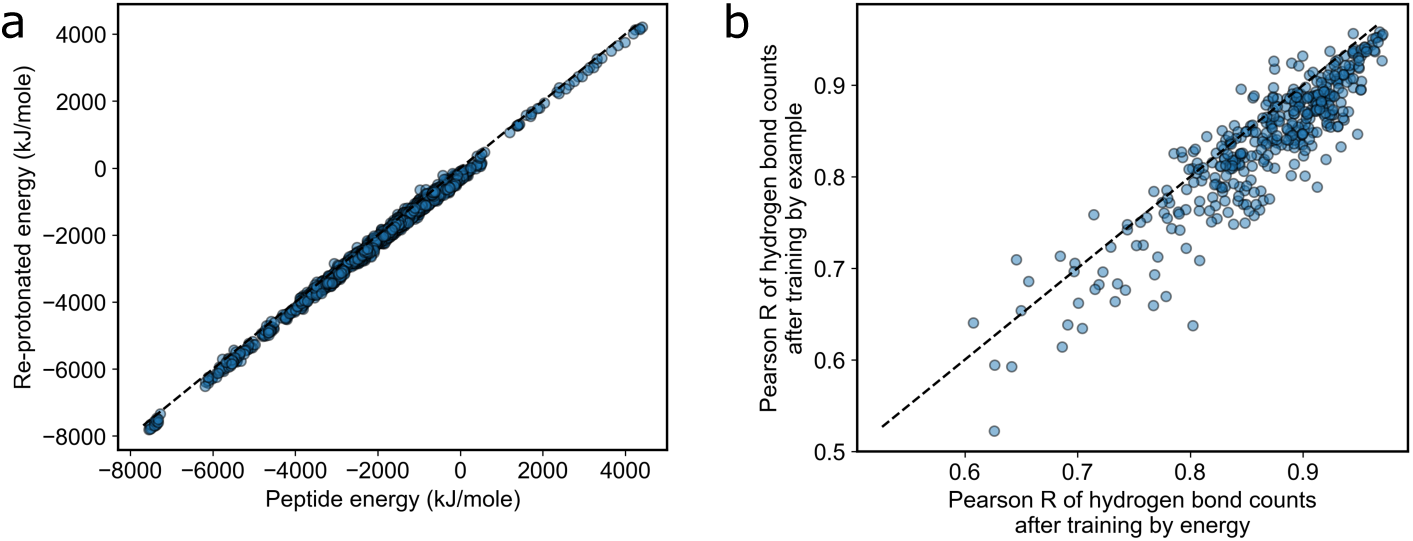
Comparison of conformations from MD simulations with re-protonated conformations after training by energy. a. Comparison of the estimated energy of conformations from MD simulations with re-protonated conformations. b. Pearson correlations between hydrogen bond counts in MD simulations and re-protonated conformations before and after training by energy.

**Supplementary Figure 12:**
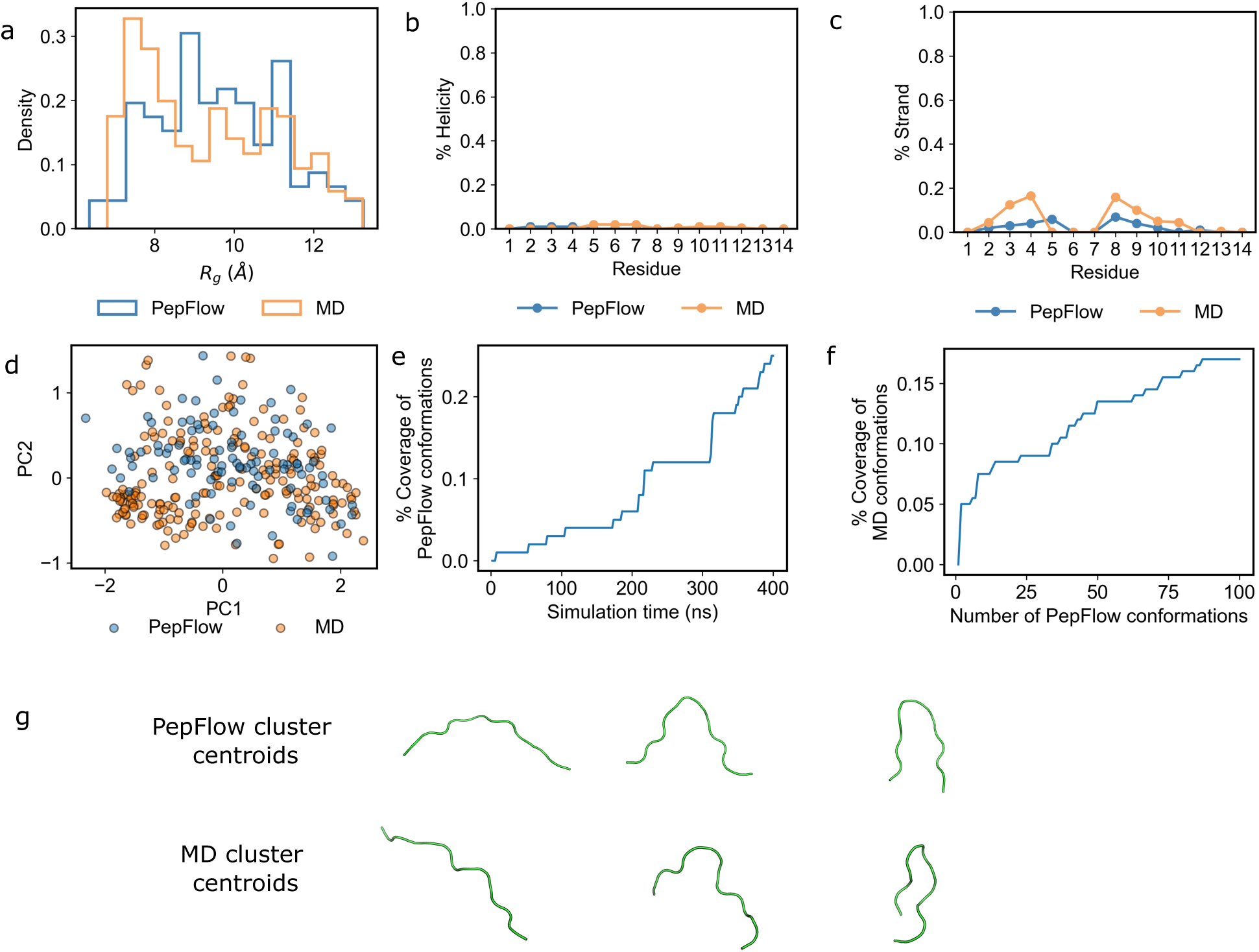
Comparison of MD and PepFlow-generated conformations for the peptide AMRLTGNKPCLYGT. a. 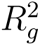 distributions from MD and PepFlow-generated conformations. b. Percentage helix formation at each residue. c. Percentage strand formation at each residue. d. Coordinate projections to principal components fit to MD conformations. e. Coverage of PepFlow-generated conformations across simulation time. f. Coverage of MD conformations as more conformations are sampled with PepFlow. g. Centroid conformations from clustering at 5 Å.

**Supplementary Figure 13:**
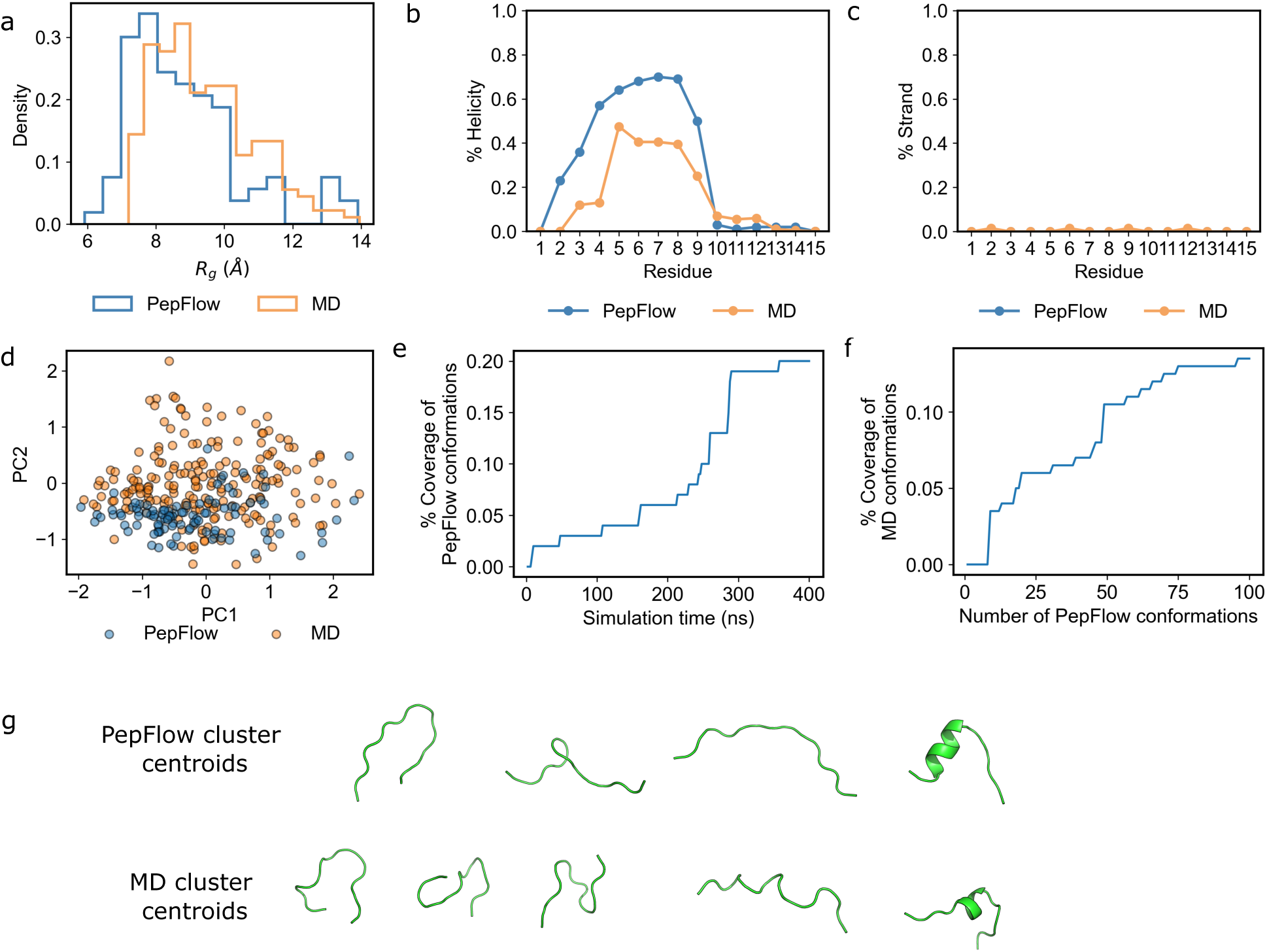
Comparison of MD and PepFlow-generated conformations for the peptide LQTKLKKLLGLESVF. a. 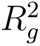 distributions from MD and PepFlow-generated conformations. b. Percentage helix formation at each residue. c. Percentage strand formation at each residue. d. Coordinate projections to principal components fit to MD conformations. e. Coverage of PepFlow-generated conformations across simulation time. f. Coverage of MD conformations as more conformations are sampled with PepFlow. g. Centroid conformations from clustering at 5 Å.

**Supplementary Figure 14:**
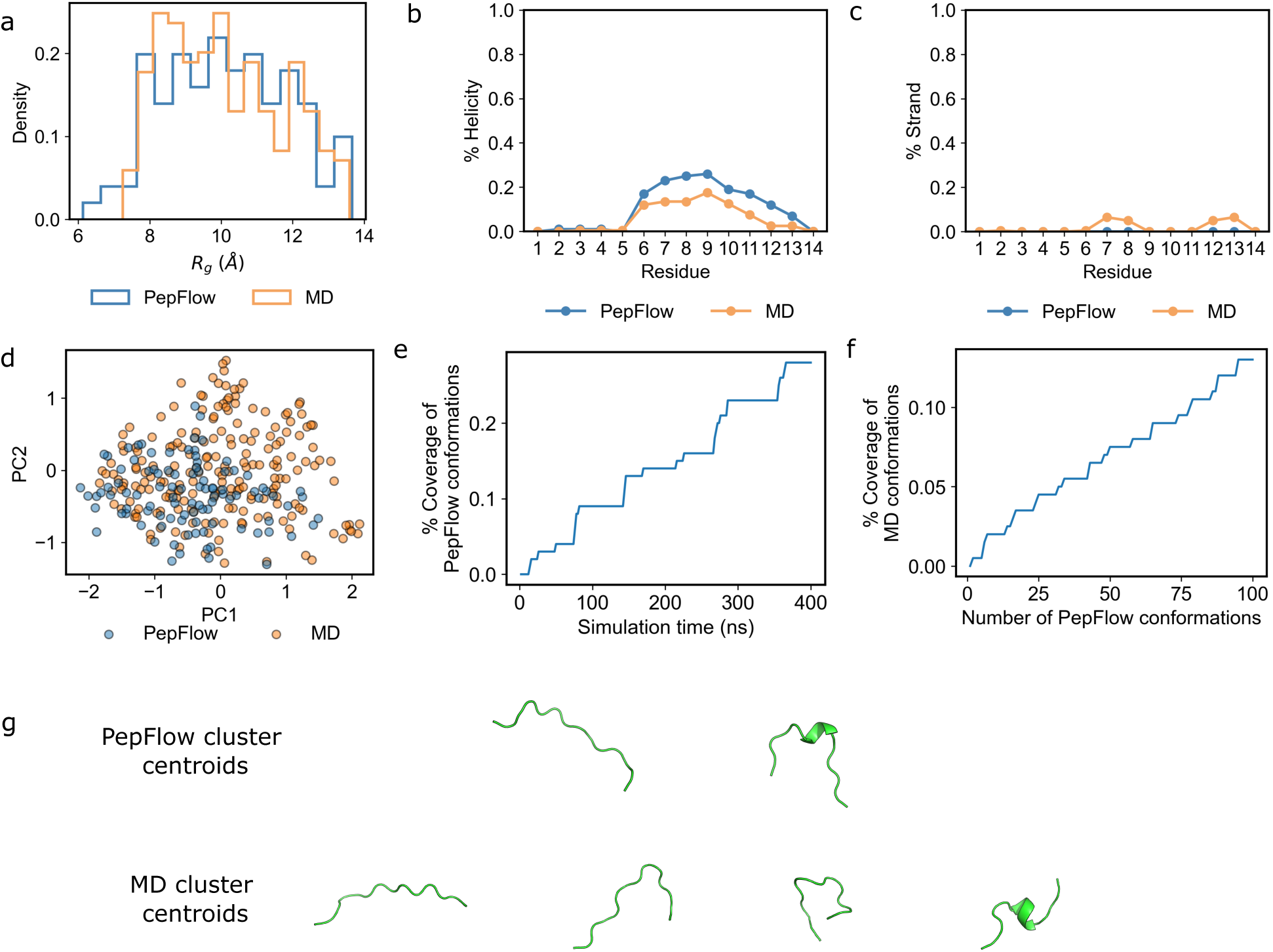
Comparison of MD and PepFlow-generated conformations for the peptide LRLKSIVSYAKKVL. a. 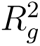 distributions from MD and PepFlow-generated conformations. b. Percentage helix formation at each residue. c. Percentage strand formation at each residue. d. Coordinate projections to principal components fit to MD conformations. e. Coverage of PepFlow-generated conformations across simulation time. f. Coverage of MD conformations as more conformations are sampled with PepFlow. g. Centroid conformations from clustering at 5 Å.

**Supplementary Figure 15:**
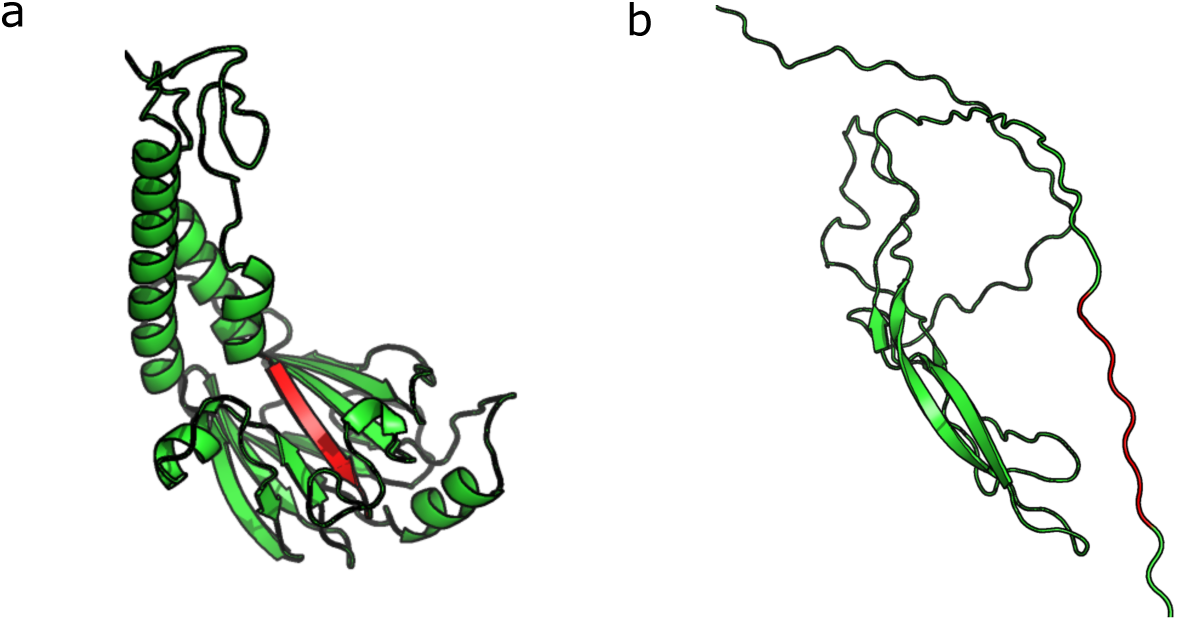
Examples of annotated SLiMs (red) in PED models. a. Annotated SLiM in PED00193. b. Annotated SLiM in PED00217.

**Supplementary Figure 16:**
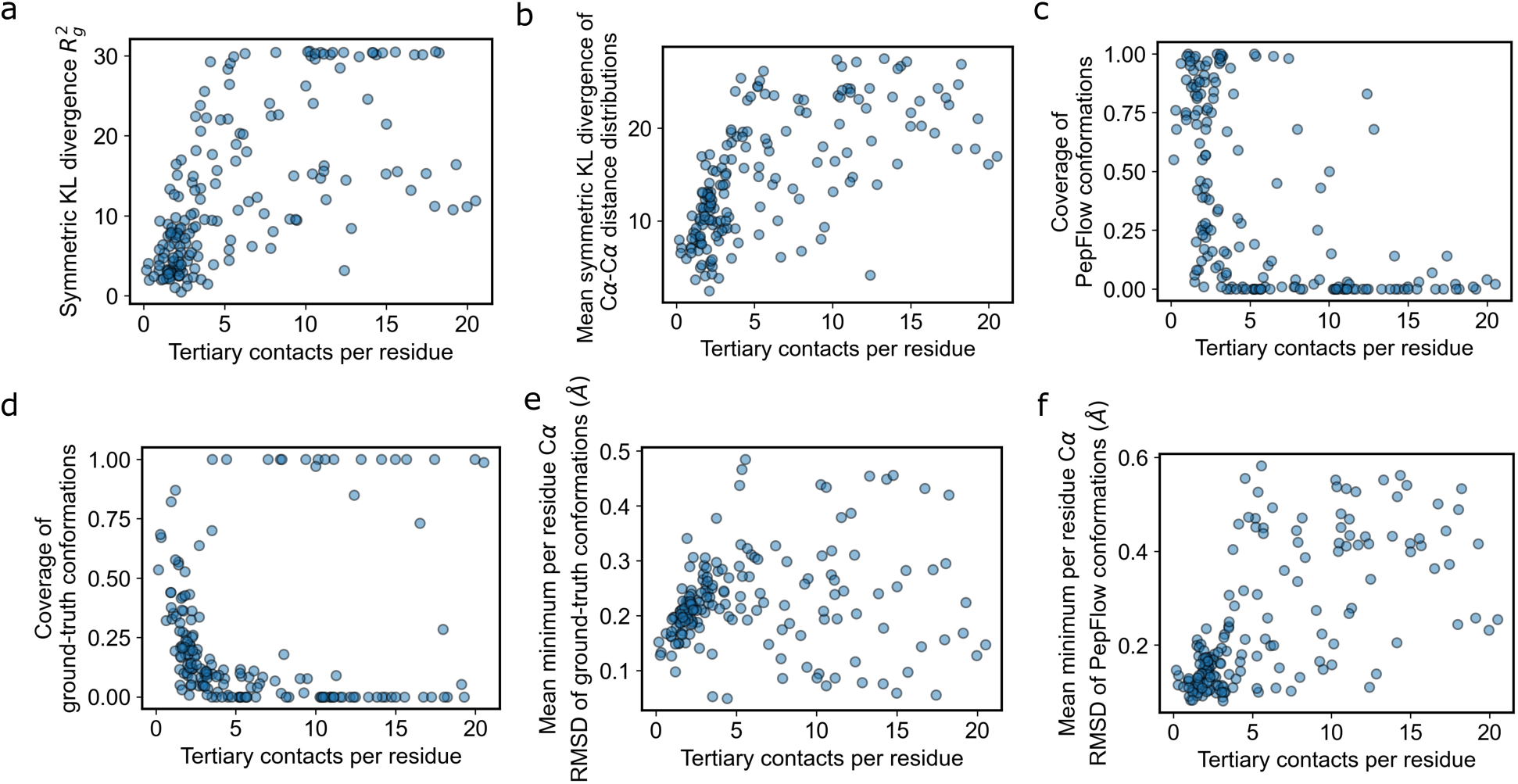
Relationship between fragment tertiary contacts and a. symmetric KL divergence of 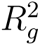 distributions b. Mean symmetric KL divergence between pairwise C*α*-C*α* distance distributions c. Coverage of PepFlow conformations at a threshold of 0.15 Å d. Coverage of SLiM conformations at a threshold of 0.15 Å e. Average minimum RMSD of SLiM conformations to PepFlow conformations f. Average minimum RMSD of PepFlow conformations to SLiM conformations.

**Supplementary Figure 17:**
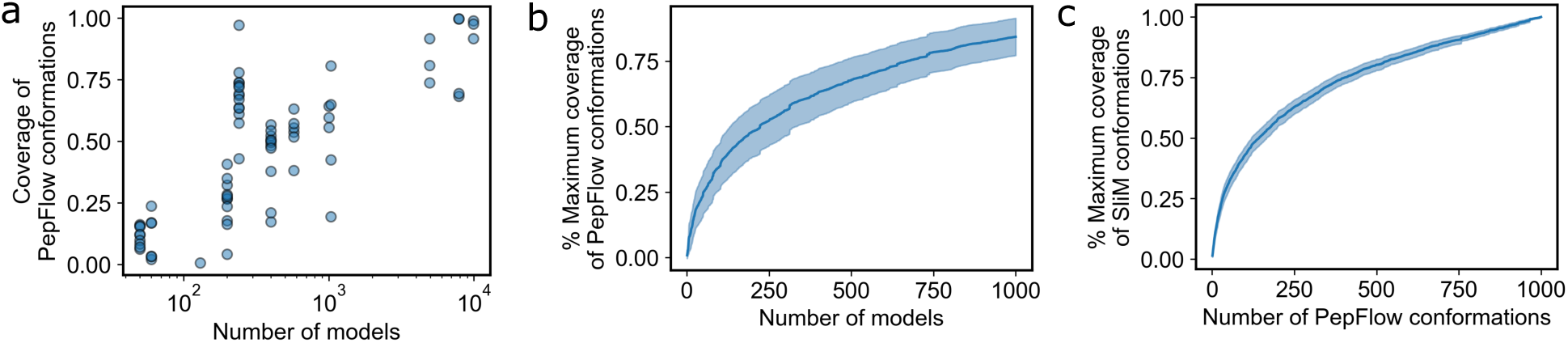
Coverage of SLiM conformations and PepFlow conformations with more sampled. a. Coverage of PepFlow conformations across SLiMs with varying structural models. b. Coverage of PepFlow-generated conformations with more SLiM conformations. c. Coverage of SLiM conformations as more conformations are sampled with PepFlow.

**Supplementary Figure 18:**
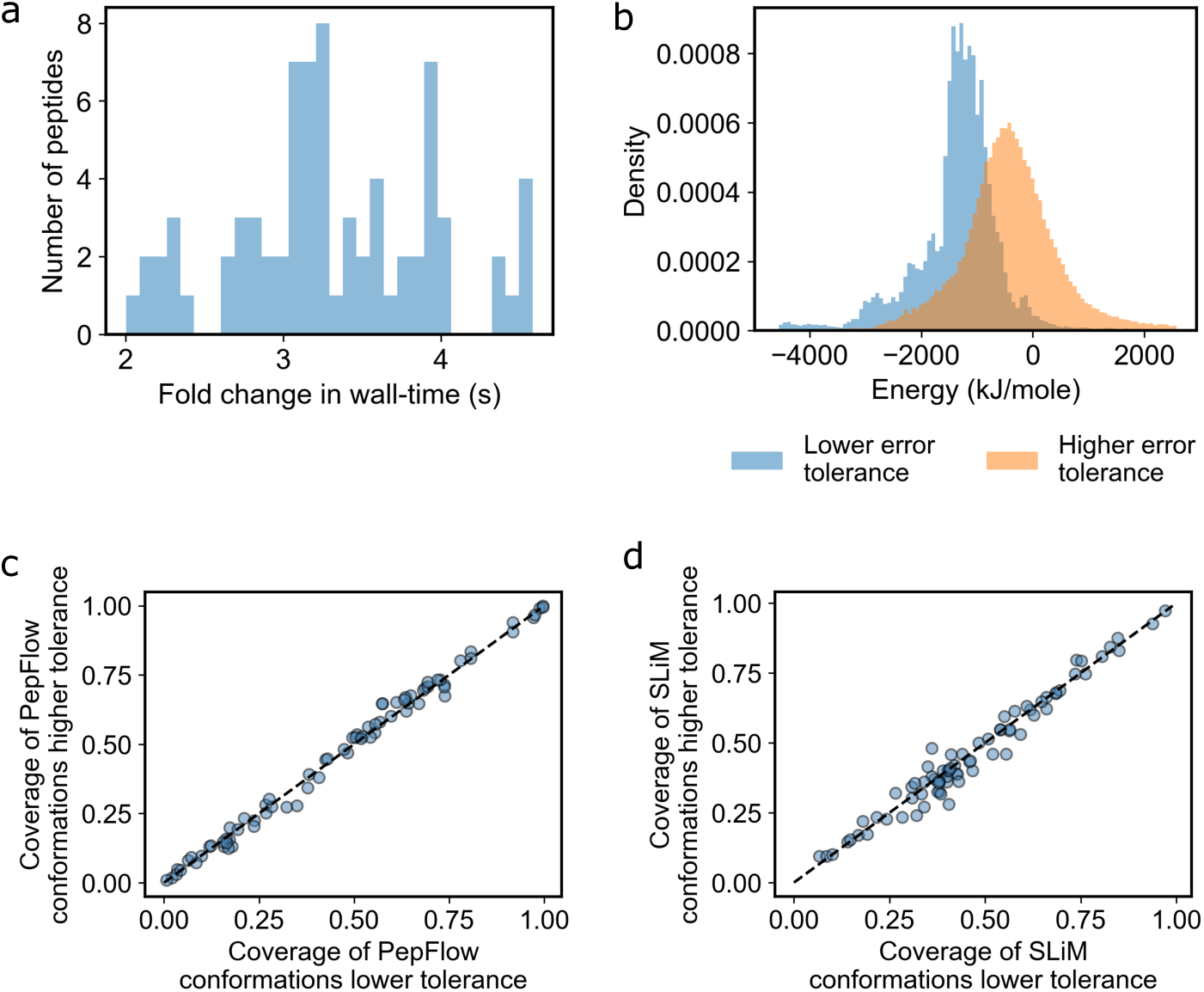
Comparison of generated conformations at low and high solver error tolerances. a. Fold change in wall-time when sampling with lower error tolerance. b. Predicted energy of conformations with low and high error tolerances. c. Comparison of coverage of PepFlow conformations by experimental SLiM conformations with different solver error tolerances. d. Comparison of coverage of experimental SLiM conformations by PepFlow-generated conformations with different solver error tolerances.

**Supplementary Figure 19:**
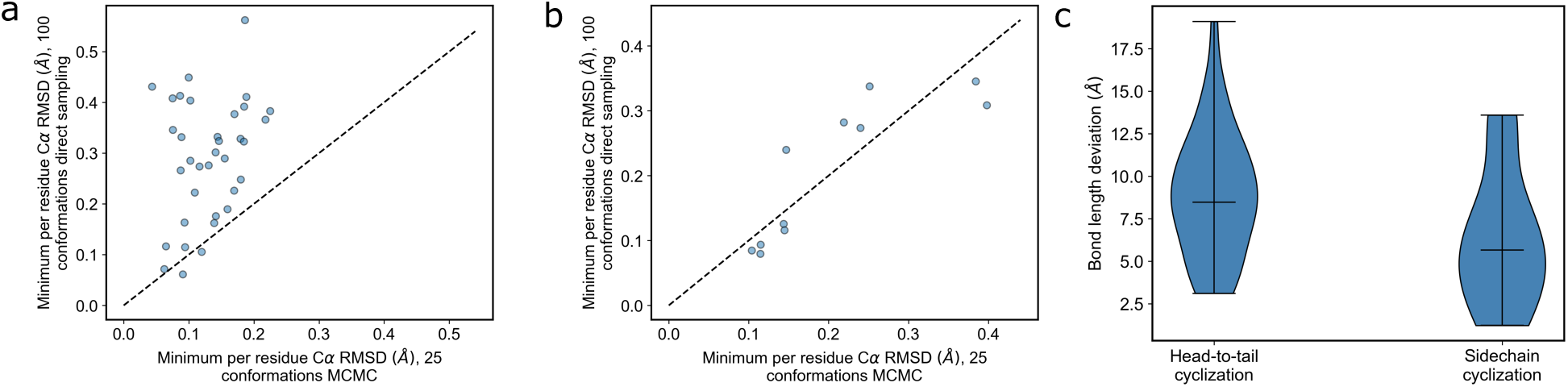
Comparison of direct sampling and MCMC for cyclic peptide structure prediction. a. Comparison of minimum RMSDs for head-to-tail cyclized peptides b. Comparison of minimum RMSD for side chain cyclized peptides. c. Minimum bond length deviations using direct sampling.

**Supplementary Figure 20:**
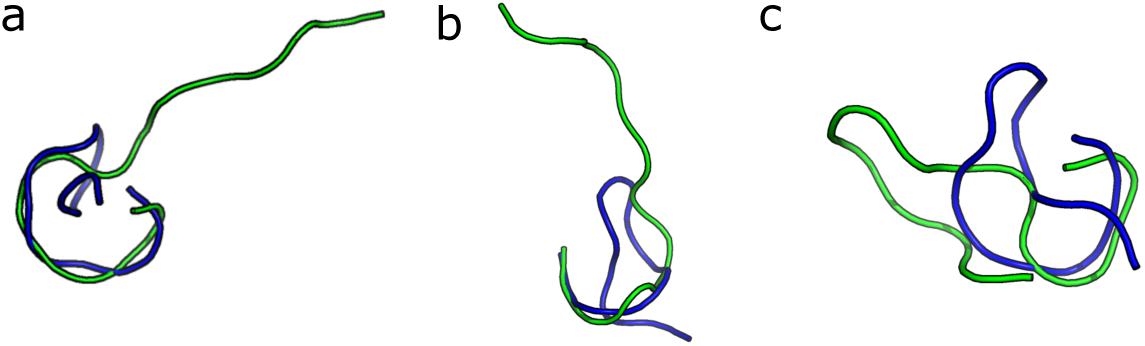
PepFlow predictions (green) compared to ground-truth lasso peptides (blue). a. Peptide LLGRSGNDRLILSKN, PDB code 7JS6. b. Peptide GG-PLAGEEMGGITT, PDB code 2MFV. c. GFGSKPLDSFGLNFF, PDB code 2N5C.

**Supplementary Figure 21:**
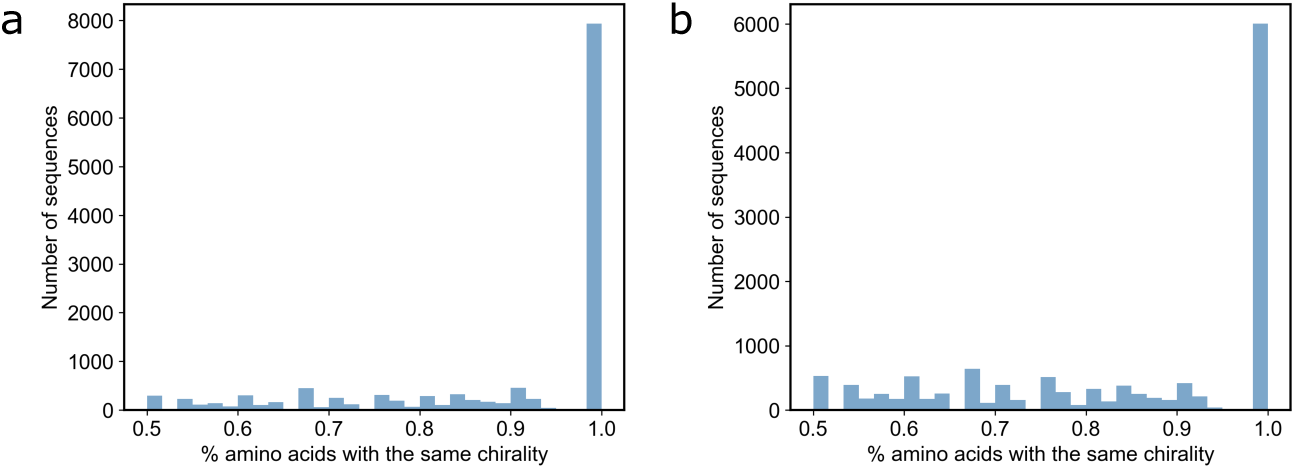
Consistency of residue chirality in generated samples. a. Distribution of the percentage of amino acids with the same chirality across different generated samples before training by energy. b. Distribution of the percentage of amino acids with the same chirality across different generated samples after training by energy.

## Notes

https://gitlab.com/oabdin/pepflow

### Competing Interest Statement

P.M.K. is a cofounder and consultant to multiple companies, including Oracle Therapeutics, TBG Therapeutics, and Zymedi. O.A. consults for Oracle Therapeutics.

